# Dihydropyrimidines sustain aggressive cancer states by stabilizing DPYSL2

**DOI:** 10.64898/2026.06.14.731825

**Authors:** Arata Hayashi, Sarah Knapp-Goldin, Abdelrahman Karmi, Vaivhav Gupta, Itai Mizrahi, Shirel Lavi, Sinai Shammah, Yael Sitton, Ravit Glukhman, Balakrishnan Solaimuthu, Sayanika Banerjee, Areej Abu Rmaileh, Anees Khatib, Michal Lichtenstein, Ora Schueler-Furman, Reuven Wiener, Yoav D. Shaul

## Abstract

Cancer cell states are governed by coordinated signaling and metabolic networks, yet the mechanisms coupling metabolic activity to oncogenic signaling remain poorly understood. Here, we identify dihydropyrimidines (DHPs), metabolites generated during pyrimidine catabolism, as “signaling metabolites” that sustain aggressive cancer cell states. Depletion of DHPs, either by knockout of the pyrimidine catabolic enzyme DPYD or by expression of the DHP-degrading enzyme DPYS, suppressed STAT3 signaling and attenuated mesenchymal and inflammatory transcriptional programs. Mechanistically, DHPs stabilized DPYSL2, a JAK1-interacting adaptor protein required for efficient STAT3 activation. Structural modeling and thermal stability analyses further support a direct interaction between DHPs and DPYSL2. Accordingly, high DPYD expression was associated with EMT, inflammatory signaling, and poor-prognosis breast cancer subtypes across patient cohorts. Ultimately, our findings establish pyrimidine catabolism as a direct regulator of signaling competence and identify DHPs as signaling-active metabolites that couple cellular metabolic state to oncogenic transcriptional programs and mesenchymal identity.

## Introduction

Cancer remains a leading cause of mortality worldwide, largely due to metastasis ^1^, which remains refractory to most therapeutic interventions despite major advances in targeted therapies and immunotherapy ^2,3^. A central contributor to these aggressive behaviors is epithelial-mesenchymal transition (EMT), a cellular program in which carcinoma cells transition from epithelial to partially mesenchymal states ^4^. During EMT, cells lose epithelial features, such as cell-cell adhesion and polarity, while acquiring mesenchymal traits that promote migration, invasion, metastatic dissemination, immune evasion, stem-like properties, and therapy resistance ^5,6^. EMT is driven by signaling pathways, such as transforming growth factor beta (TGFβ) ^7^, yet the mechanisms that stabilize aggressive cell states along the EMT spectrum remain incompletely understood. One signaling pathway strongly linked to aggressive mesenchymal states is the interleukin-6 (IL-6)-Janus kinase (JAK)-signal transducer and activator of transcription 3 (STAT3) axis ^8,9^. Aberrant IL6-JAK-STAT3 signaling promotes cancer cell survival, invasion, immune modulation, metastasis, and cancer stem-like phenotypes ^10,11^. However, the mechanisms that sustain STAT3 activity and couple inflammatory signaling to aggressive cancer cell states remain poorly defined. A defining feature of malignant transformation is the profound reprogramming of cellular metabolism ^12^. Beyond supporting anabolic growth ^13–15^, metabolic reprogramming actively shapes tumor behavior by regulating stress adaptation, immune interactions, and cellular plasticity ^10,11,16^. Increasing evidence indicates that these noncanonical functions of metabolic processes are mediated not only by enzymes but also by metabolites that directly govern the epigenetic landscape and signal transduction pathways ^17^. These signaling functions can occur through covalent mechanisms, including metabolite-dependent chromatin and protein modifications ^18^, as well as non-covalent interactions that regulate protein activity and stability ^19,20^. Classic examples include oncometabolites produced by mutant isocitrate dehydrogenases (IDH1 and IDH2) ^21^ and nutrient-sensitive metabolites that reshape transcriptional programs, including the mTOR pathway ^22^, stress responses ^23^, immune interactions ^24^, and cell fate decisions ^10,11,16^. Metabolic profiling studies have demonstrated that EMT is accompanied by extensive metabolic remodeling ^25^, while accumulation of the classical oncometabolite fumarate following fumarate hydratase (FH) loss directly promotes EMT activation and aggressive cellular phenotypes ^26,27^. However, most studies of metabolic-signaling paradigms described to date have focused on central carbon metabolism, amino acid metabolism, or lipid-derived signals ^20^. In contrast, the signaling potential of nucleotide catabolic intermediates has been largely unexplored.

In cancer, pyrimidine metabolism has been studied predominantly in the context of nucleotide biosynthesis ^28,29^, whereas pyrimidine catabolism remains comparatively unexplored. This metabolic pathway is initiated with dihydropyrimidine dehydrogenase (DPYD), which catalyzes the rate-limiting step of pyrimidine degradation, reducing uracil and thymine to their corresponding dihydropyrimidines, 5,6-dihydrouracil (DHU) and 5,6-dihydrothymine (DHT), collectively referred to as dihydropyrimidines (DHPs) ^30^. DHPs are subsequently processed by the downstream enzymes dihydropyrimidinase (DPYS) and β-ureidopropionase ^30^. This pathway is known for hepatic pyrimidine turnover and for the inactivation of the chemotherapeutic agent 5-fluorouracil (5-FU) ^31^, where DPYD loss-of-function variants are major determinants of fluoropyrimidine toxicity ^32^. However, emerging evidence suggests that pyrimidine catabolism also has non-canonical functions in aggressive cancer cell states. We previously showed that the EMT program selectively expresses the upstream enzyme DPYD while lacking DPYS, resulting in the accumulation of DHPs, which is essential for the proper execution of this program ^16^. In parallel, independent work demonstrated that thymidylate synthase (TYMS) maintains the de-differentiated state of triple-negative breast cancer by fueling DPYD-dependent pyrimidine catabolism ^33^. Together, these findings point towards a non-canonical role for DHPs as signaling-regulatory metabolites that couple pyrimidine catabolism to oncogenic pathway activity and the acquisition of aggressive cancer cell states.

Here, we reveal that intracellular DHPs act as signaling-regulatory metabolites that control the stability of dihydropyrimidinase-like 2 (DPYSL2/CRMP2), a structural homolog of DPYS that lacks enzymatic activity and serves as a critical scaffold for JAK1-STAT3 signaling ^34^. We show that DHP abundance, through a post-translational mechanism, stabilizes DPYSL2, thereby sustaining STAT3 pathway activation. Structural modeling identified a pseudoenzyme DHP-binding pocket in DPYSL2, providing a potential structural basis for metabolite-dependent stabilization. Consistently, depletion of DHPs through DPYD loss or ectopic DPYS expression destabilizes DPYSL2 and suppresses mesenchymal transcriptional programs. Together, these findings establish pyrimidine catabolism as a metabolically encoded regulator of oncogenic signaling and identify the DHP-DPYSL2-STAT3 axis as a mechanism coupling nucleotide catabolism to aggressive mesenchymal cancer cell states.

## Results

### DPYD expression correlates with mesenchymal cell states and STAT3 signaling across cancers

To investigate whether pyrimidine catabolism is linked to oncogenic signaling programs, we analyzed public transcriptomic datasets using Gene ENrichment Identifier (GENI) platform ^35^. DPYD expression strongly associates with the Hallmarks of epithelial-to-mesenchymal transition (EMT), inflammatory signaling, and the IL6-JAK-STAT3 pathway ^36^ across multiple cancer types, including breast, lung, pancreatic, and glioblastoma (Fig. 1A, B, and Fig. S1A, B). We next investigated the relationship between DPYD expression and STAT3 pathway activity, as this signaling cascade plays a central role in driving EMT and tumor aggressiveness ^10,11,34^. In breast cancer cohorts, DPYD expression positively correlated with both interleukin 6 (*IL6)* and Suppressor of cytokine signaling 3 (*SOCS3*), a canonical STAT3 target gene (Fig. 1C) ^37^. Similarly, induction of EMT in A549 lung adenocarcinoma cells by TGFβ was accompanied by progressive upregulation of DPYD together with increased N-cadherin and reduced E-cadherin expression (Fig. S1C). This pattern was further validated at the protein level, with basal/mesenchymal breast cancer cell lines exhibiting markedly higher DPYD abundance than luminal/epithelial cell lines (Fig. S1D). Interestingly, while DPYD was also highly expressed in mouse liver tissue, mesenchymal markers remained restricted to basal cancer cell lines, suggesting that DPYD’s canonical metabolic role is context dependent (Fig. S1E).

**Figure 1.**
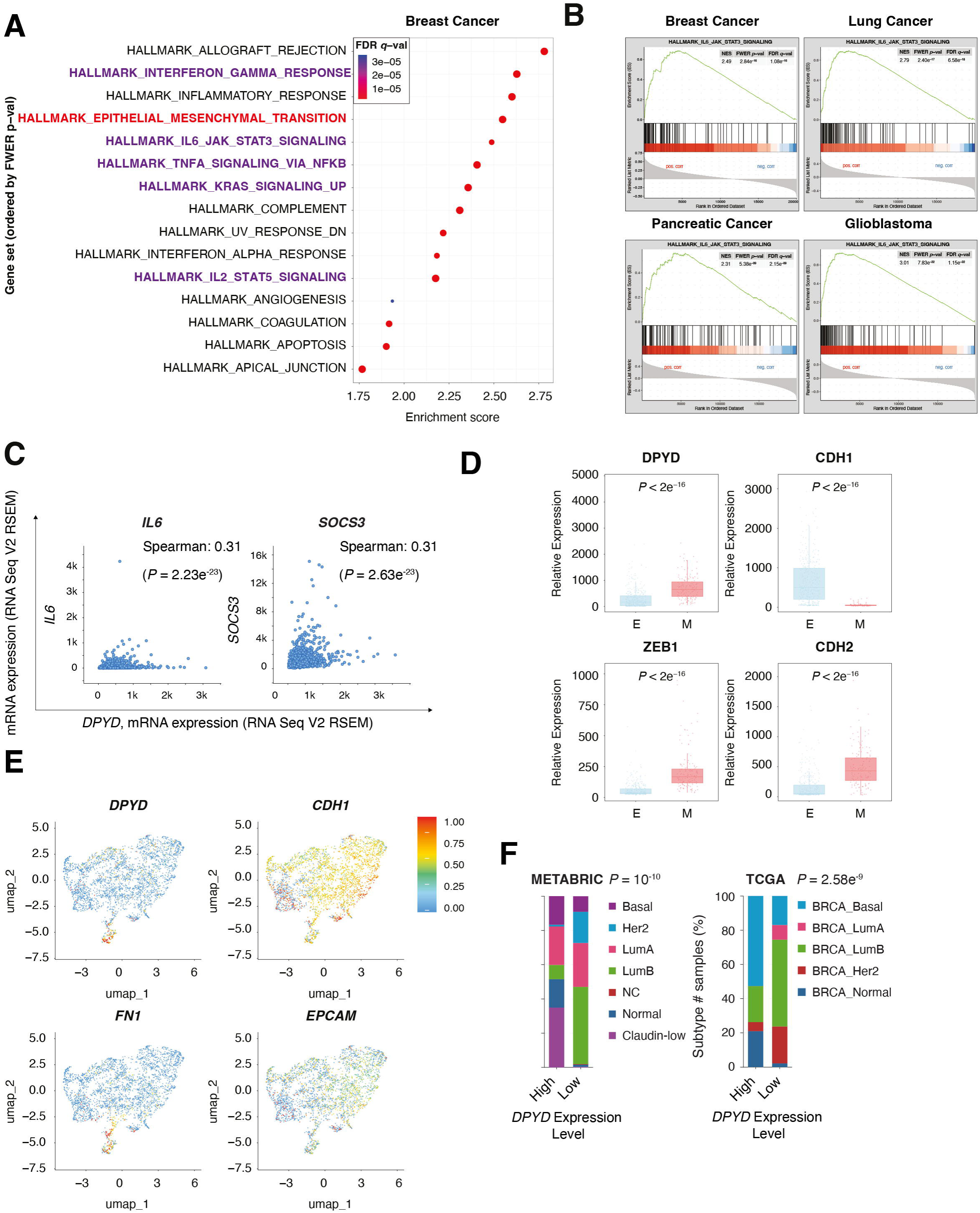
DPYD expression converges with mesenchymal cell states and IL6-JAK-STAT3 signaling across cancers. **(A)** DPYD expression is associated with EMT and inflammatory signaling pathways in breast cancer. Gene set enrichment analysis using the GENI platform showing Hallmark pathways positively associated with *DPYD* expression in breast cancer (TCGA, Firehose Legacy). Pathways are ranked by FWER *P* value, and point color indicates FDR *q* value. EMT-, inflammatory-, and oncogenic signaling-related pathways are highlighted. Statistical significance was assessed using the GENI GSEA workflow, with FWER *P* values and FDR *q* values shown. **(B)** DPYD expression correlates with IL6-JAK-STAT3 signaling across multiple cancer types. GSEA plots showing enrichment of the Hallmark IL6-JAK-STAT3 signaling pathway in gene expression profiles ranked by their correlation with DPYD across breast, lung, pancreatic cancer, and glioblastoma datasets (TCGA, Firehose Legacy). Normalized enrichment scores (NES), FWER *P* values, and FDR *q* values are indicated. **(C)** DPYD expression positively correlates with the STAT3 targets *IL6* and *SOCS3* in breast cancer patients. Correlation analysis of *DPYD* mRNA expression with *IL6* and *SOCS3* expression in breast cancer datasets. Spearman correlation coefficients and *P* values are indicated and were calculated by the cBioPortal. **(D)** DPYD expression is elevated in mesenchymal-like breast cancer cell lines. Cancer cell lines were divided into epithelial (E, *n* = 378 cell lines) and mesenchymal (M, *n* = 150 cell lines) groups based on the expression of known mesenchymal markers using the MERAV platform. Box-and-whisker plots with overlaid individual data points show the relative expression levels of *DPYD*, the epithelial marker *CDH1*, and the mesenchymal markers *ZEB1* and *CDH2* in each group. The *P* value was determined by the Wilcoxon rank-sum test. **(E)** DPYD expression marks mesenchymal-like cells at the single-cell level. UMAP visualization of breast cancer single-cell RNA-seq data showing expression patterns of *DPYD,* the mesenchymal marker *FN1*, and epithelial markers *CDH1* and *EPCAM*. Color scale indicates normalized relative expression levels, scaled between 0 and 1. **(F)** DPYD expression is elevated in aggressive breast cancer subtypes. Breast cancer samples were divided into DPYD-high and DPYD-low groups (≥1 s.d. above or below the mean DPYD expression, respectively). For each group, the percentage of breast cancer subtypes is color-coded. Breast cancer data were obtained from the METABRIC (left) or TCGA (PanCancer Atlas project; right) databases, and the *P* values were calculated by cBioPortal. LumA, luminal A; LumB, luminal B.

Consistent with a mesenchymal cell-state association, analysis of the MERAV dataset ^38^ demonstrated that DPYD expression is significantly elevated in mesenchymal compared with epithelial breast cancer cell lines (Fig. 1D). To extend these observations to the single-cell level, we analyzed publicly available breast cancer single-cell RNA-seq data. DPYD expression closely mirrored the distribution of the mesenchymal marker *FN1*, while remaining largely mutually exclusive with the epithelial markers *CDH1* and *EPCAM* (Fig. 1E).

Finally, analysis of the METABRIC ^39,40^ and TCGA PanCancer Atlas breast cancer cohorts ^41^, available on the cBioPortal ^42^, revealed that tumors with high DPYD expression are significantly enriched in claudin-low and basal subtypes (Fig. 1F). Together, these findings identify DPYD as, a marker of mesenchymal cell states and aggressive breast cancer subtypes, closely linked to the IL6-JAK-STAT3 axis across multiple systems and patient cohorts.

### Intracellular dihydropyrimidine abundance governs STAT3 signaling

Upon establishing a robust association between DPYD expression and the IL6-JAK-STAT3 signaling axis across cancers, we next asked whether intracellular DHP abundance directly regulates STAT3 pathway activity. To reduce intracellular DHP levels, we either knocked out the DHP-generating enzyme DPYD (DPYD-KO) or ectopically expressed the DHP-catabolizing enzyme DPYS-FLAG in the basal B breast cancer cell line MDA-MB-231 (Fig. 2A and B). As an enzymatic control, we generated a catalytically inactive DPYS mutant (DPYS-K159A-FLAG). Lysine 159 (K159) was selected based on UniProt annotations and sequence alignment of human DPYS with structurally characterized DPYS homologs from *Saccharomyces cerevisiae* and *Dictyostelium* ^43,44^. This analysis identified K159 as a highly conserved residue located adjacent to predicted catalytic and substrate-binding regions (Fig. S2A). Accordingly, an *in vitro* DPYS enzymatic assay ^45^ further confirmed that wild-type DPYS, but not the K159 mutant (K159L), efficiently catalyzed DHU turnover (Fig. S2B). We focused on DHU because targeted metabolomics indicated that it is the predominant DHP species in MDA-MB-231 cells (Fig. S2C). Consistent with this enzymatic assay, both DPYD loss and wild-type DPYS overexpression significantly reduced intracellular DHU levels, whereas expression of the DPYS-K159A mutant failed to alter DHU abundance (Fig. 2C), independently validating K159 as a previously uncharacterized residue essential for DPYS catalytic activity. Importantly, these manipulations did not significantly affect cell proliferation, excluding the possibility of altered growth kinetics (Fig. S2D).

**Figure 2.**
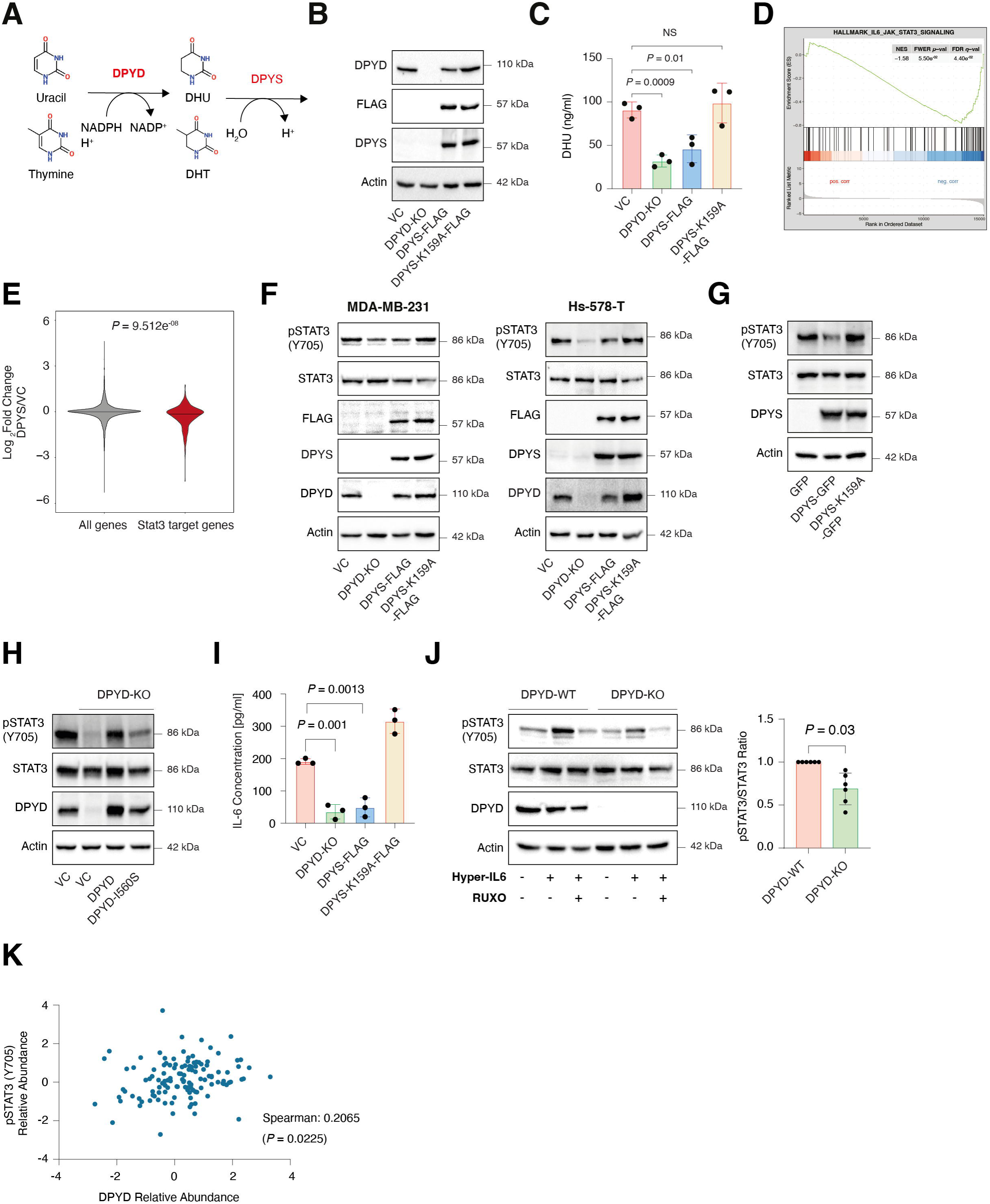
Intracellular dihydropyrimidine abundance governs STAT3 signaling output. **(A)** DPYD and DPYS control intracellular DHP abundance. Schematic representation of the pyrimidine degradation pathway. DPYD reduces uracil and thymine to the dihydropyrimidines DHU and DHT, which are subsequently processed by DPYS. **(B)** DPYD loss and DPYS overexpression establish genetic tools to reduce DHP abundance. Immunoblots representing vector control (VC), DPYD knockout (DPYD-KO), DPYS-FLAG overexpression, and DPYS-K159A-FLAG expression in MDA-MB-231 cancer cell lines. Cells were lysed and subjected to immunoblotting using the indicated antibodies. **(C)** DPYD loss and wild-type DPYS overexpression reduce intracellular DHU levels. Targeted metabolite quantification of intracellular DHU levels in the indicated MDA-MB-231 cell lines. Each value represents the mean ± s.d. (*n* = 3). *P* values were determined by an unpaired two-sided Student’s *t*-test. **(D)** DPYS expression suppresses the Hallmark IL6-JAK-STAT3 signaling program. MDA-MB-231 vector control and DPYS-FLAG-expressing cells were subjected to RNA-seq analysis. Genes were ranked by differential expression between DPYS-FLAG and vector control cells and then subjected to GSEA. The normalized enrichment score (NES), FWER *P* value, and FDR *q* value were computed by GSEA. **(E)** DPYS expression suppresses STAT3 target-gene expression. Violin plot showing the distribution of log2 fold-change values for all genes compared with DoRothEA-defined STAT3 target genes following DPYS overexpression. The *P* value was determined by the Wilcoxon rank-sum test. **(F)** STAT3 phosphorylation correlates with DHP intracellular abundance. Immunoblots representing MDA-MB-231 cells (left) and Hs-578-T cells (right) expressing vector control (VC), DPYD knockout, DPYS-FLAG, or DPYS-K159A-FLAG. Cells were lysed and subjected to immunoblotting using the indicated antibodies. Total STAT3 and actin are shown as controls. **(G)** Inducible DPYS expression suppresses STAT3 phosphorylation in a catalytic activity-dependent manner. MDA-MB-231 cells expressing doxycycline-inducible DPYS-GFP or DPYS-K159A-GFP were treated with 2 μg/ml doxycycline for 96 h, lysed, and subjected to immunoblotting using the indicated antibodies. **(H)** DPYD enzymatic activity is required to restore STAT3 phosphorylation. Immunoblot analysis of DPYD-knockout cells reconstituted with wild-type human DPYD or the catalytically attenuated DPYD-I560S mutant. Re-expression of wild-type DPYD restores STAT3 phosphorylation, whereas DPYD-I560S fails to rescue pathway activation. **(I)** DHP depletion reduces IL-6 secretion. The indicated MDA-MB-231 cell lines were cultured for 48 h, after which conditioned media were collected, and IL-6 levels were measured using a specific ELISA kit. Each value represents the mean ± s.d. (*n* = 3). **(J)** DPYD loss attenuates Hyper-IL-6-induced STAT3 activation. DPYD-WT and DPYD-KO MDA-MB-231 cells were treated with 5 pg/ml Hyper-IL-6 for 1 h, with or without pretreatment with 1 μM ruxolitinib (RUXO). Left: cells were lysed and subjected to immunoblotting using the indicated antibodies. Right: quantification of phospho-STAT3 immunoblots normalized to total STAT3 levels. Each bar represents the mean ± s.d. (*n* = 6). The *P* value was determined by a one-sample Wilcoxon signed-rank test. **(K)** DPYD protein abundance positively correlates with STAT3 Y705 phosphorylation in breast cancer. Correlation analysis of DPYD relative abundance and STAT3 pY705 relative abundance in CPTAC breast cancer samples. Spearman correlation coefficients and *P* values are indicated and were calculated by the cBioPortal.

We next asked whether this experimentally induced reduction in intracellular DHP abundance impacts STAT3 signaling output. RNA sequencing of MDA-MB-231 cells expressing either empty vector or DPYS-FLAG revealed a major global transcriptional shift upon DHP depletion (Fig. S2E, S2F, and Supplementary Table 1). Gene set enrichment analysis (GSEA) ^46^ showed that the “Hallmark IL6-JAK-STAT3 signaling” program was among the most significantly downregulated pathways (Fig. 2D, Fig. S2G and S2H), and DoRothEA analysis ^47^ demonstrated significant suppression of the STAT3 regulon (Fig. 2E and S2I). Among the downregulated genes were multiple inflammatory and STAT3-associated factors, including *IL6*, *IL1B*, *TNF*, *VEGFA*, and *ICAM1*, whose reduced expression was further validated by quantitative RT-PCR (Fig. S2J). At the signaling level, we identified that STAT3 phosphorylation at Y705 (pSTAT3), a well-established marker of STAT3 pathway activation ^37^, closely tracked intracellular DHP abundance. Specifically, pSTAT3 was reduced in both DPYD-deficient and DPYS-overexpressing MDA-MB-231 and Hs-578-T cells, while remaining largely preserved in cells expressing the catalytically impaired DPYS-K159A mutant (Fig. 2F). Additionally, we found that induction of wild-type DPYS for 96 h reduced STAT3 phosphorylation, in contrast to the DPYS-K159A mutant, confirming that DPYS catalytic activity is required for suppression of STAT3 signaling (Fig. 2G and S2K). Furthermore, to directly determine whether DPYD catalytic activity is required for this signaling axis, we reconstituted DPYD-knockout cells with either wild-type human DPYD or a catalytically attenuated mutant (I560S) ^16^. Whereas introducing wild-type DPYD robustly restored STAT3 phosphorylation, the catalytic mutant failed to rescue STAT3 phosphorylation (Fig. 2H). Consistently, IL-6 secretion, a readout for STAT3 activity ^11^, was significantly and positively correlated with DHP levels (Fig. 2I).

Finally, to determine whether DPYD is required for STAT3 activation downstream of cytokine receptor signaling, we stimulated WT and DPYD-deficient cells with Hyper-IL-6, which directly activates the IL-6 receptor-gp130 signaling complex ^48^. While Hyper-IL-6 strongly induced STAT3 phosphorylation in control cells, DPYD loss attenuated this response, indicating that DPYD is required for efficient STAT3 pathway activation even under forced cytokine stimulation conditions (Fig. 2J). Moreover, pharmacological inhibition of JAK1 signaling by ruxolitinib abolished Hyper-IL-6-induced STAT3 phosphorylation, confirming pathway specificity. Consistent with these experimental findings, analysis of patient proteomic datasets further revealed a significant positive correlation between DPYD abundance and STAT3 Y705 phosphorylation across tumors (Fig. 2K). Together, these findings demonstrate that intracellular DHP abundance controls STAT3 signaling through a DPYD-dependent mechanism and sustains an IL6-driven autocrine feed-forward loop in aggressive breast cancer cells.

### DHPs regulate STAT3 signaling by stabilizing DPYSL2

Having established that intracellular DHP abundance sustains STAT3 signaling, we next sought to identify the molecular intermediary linking pyrimidine metabolism to pathway activation. Quantitative proteomic analysis revealed that DPYSL2 was among the most significantly downregulated proteins following DPYS overexpression (Fig. 3A and Supplementary Table 2). Immunoblot and immunofluorescence analyses independently confirmed this reduction in DPYSL2 abundance (Fig. 3B and Fig. S3A), whereas DPYSL2 mRNA expression remained unchanged (Fig. S3B). Furthermore, the abundance of ectopically expressed DPYSL2-GFP was markedly reduced in DPYD-KO cells, indicating that DPYD is required for DPYSL2 stability independently of its endogenous promoter context (Fig. 3C). Consistent with these findings, CPTAC breast cancer proteomic datasets ^4949^ revealed a significant positive correlation between DPYD and DPYSL2 protein abundance across patient tumors, supporting the clinical relevance of the DHP-DPYSL2 signaling axis (Fig. 3D). Finally, induction of wild-type DPYS reduced DPYSL2 protein abundance, whereas induction of the catalytically impaired DPYS-K159A mutant failed to destabilize DPYSL2, further establishing that DPYS catalytic activity is required for DPYSL2 destabilization (Fig. 3E). Together, these findings indicate that intracellular DHP abundance positively regulates DPYSL2 protein stability. Intriguingly, sequence alignment revealed extensive similarity between human DPYS and DPYSL2, particularly around the catalytic K159 residue of DPYS (Fig. S3C). Although DPYSL2 lacks the catalytic activity of DPYS ^5050^, its strong structural similarity to the enzyme raised the possibility that it retains metabolite-binding properties and acts as a DHP-responsive signaling protein.

**Figure 3.**
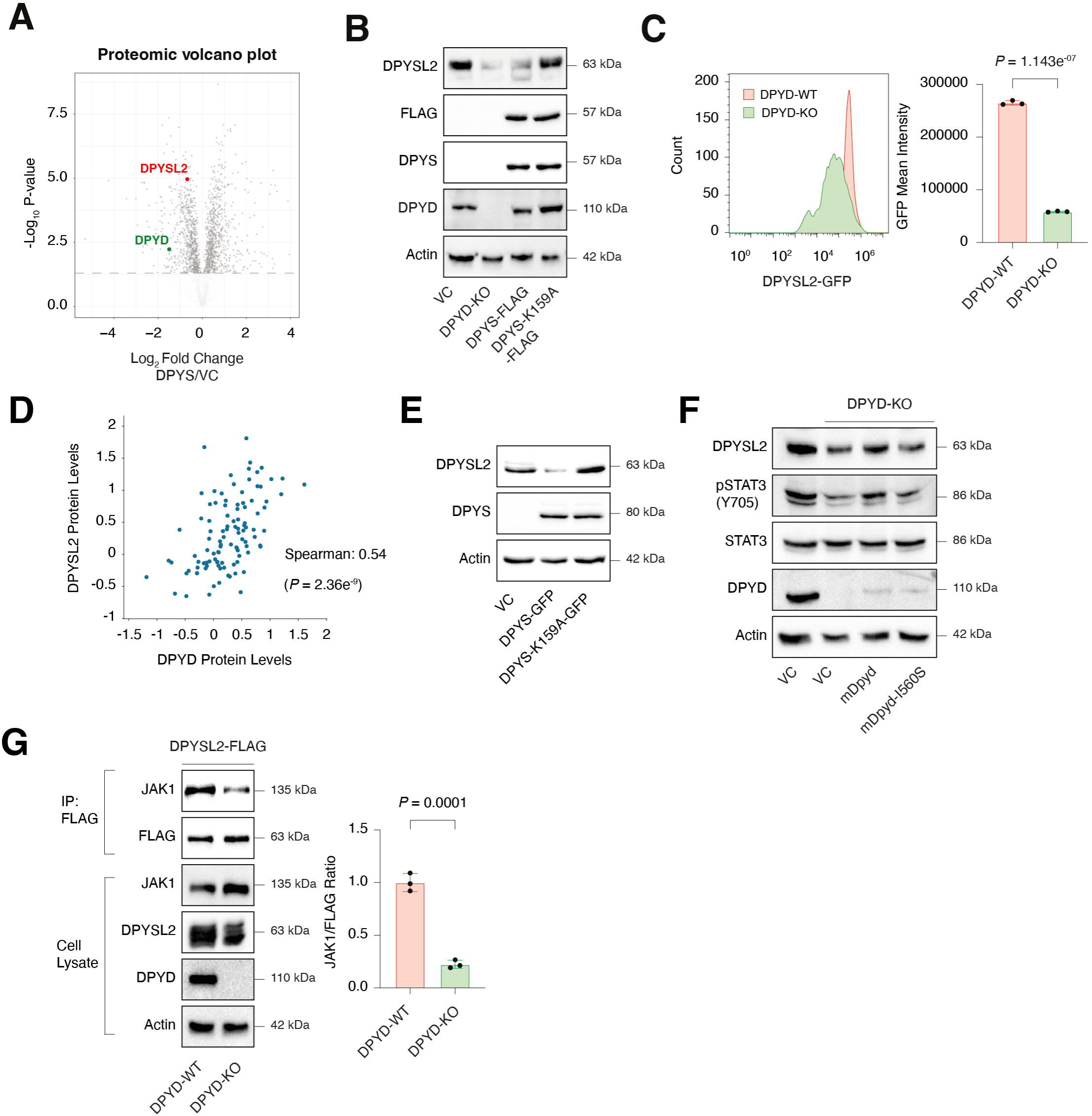
DHP abundance stabilizes DPYSL2 to sustain STAT3 signaling. **(A)** DPYS overexpression reduces DPYSL2 protein abundance. Proteomic volcano plot comparing DPYS-FLAG overexpressing MDA-MB-231 cells with vector control cells. DPYSL2 is highlighted among proteins reduced following DPYS overexpression. **(B)** DPYSL2 protein abundance depends on intracellular DHP levels. Immunoblots representing vector control (VC), DPYD knockout (DPYD-KO), DPYS-FLAG overexpression, and DPYS-K159A-FLAG expression in MDA-MB-231 cells. Cells were lysed and subjected to immunoblotting using the indicated antibodies. **(C)** DPYD loss destabilizes ectopically expressed DPYSL2-GFP. Flow cytometry analysis of DPYSL2-GFP fluorescence intensity in DPYD-WT and DPYD-KO MDA-MB-231 cells. Left: representative histogram of GFP fluorescence intensity. Right: quantification of GFP-mean intensity. Each value represents the mean ± s.d. (*n* = 3). The *P* value was determined by an unpaired two-tailed Student’s *t*-test. **(D)** DPYD and DPYSL2 protein levels positively correlate in breast cancer. Correlation analysis of DPYD and DPYSL2 protein abundance in CPTAC breast cancer samples. Spearman correlation coefficients and *P* values are indicated and were calculated by the cBioPortal. **(E)** Inducible DPYS expression reduces DPYSL2 abundance in a catalytic activity-dependent manner. MDA-MB-231 cells expressing doxycycline-inducible DPYS-GFP or DPYS-K159A-GFP were treated with doxycycline for 96 h, lysed, and subjected to immunoblotting using the indicated antibodies. **(F)** DPYD catalytic activity restores DPYSL2 and STAT3 signaling. Immunoblot analysis of DPYD-KO MDA-MB-231 cells reconstituted with wild-type murine Dpyd (mDpyd) or the catalytically attenuated mDpyd-I560S mutant. Cells were lysed and subjected to immunoblotting using the indicated antibodies. **(G)** DHP abundance regulates the DPYSL2-JAK1 interaction. Cells expressing DPYSL2-FLAG in DPYD-WT or DPYD-KO backgrounds were lysed and subjected to immunoprecipitation using anti-FLAG antibodies. Cell lysates and immunoprecipitants were analyzed by immunoblotting for the indicated proteins. Right, quantification of the JAK1/FLAG immunoblot ratio. Each bar represents the mean ± s.d. (*n* = 3). *P* values were determined by an unpaired two-sided Student’s t-test.

Since DPYSL2 is a critical regulator of JAK1-STAT3 signaling ^34^, we hypothesized that intracellular DHP regulates STAT3 activity by modulating DPYSL2 stability. We found that reducing DHP levels through DPYD loss impacts DPYSL2 and STAT3 phosphorylation levels (Fig. 3F). Importantly, reconstitution of DPYD-knockout cells with wild-type murine DPYD restored both DPYSL2 abundance and STAT3 phosphorylation, whereas the catalytically attenuated I560S mutant failed to rescue either phenotype, further demonstrating that DPYD enzymatic activity is required to sustain the DPYSL2-STAT3 signaling axis (Fig. 3F). Consistent with a post-transcriptional mechanism, wild-type murine DPYD increased DPYSL2 protein abundance without significantly altering DPYSL2 mRNA levels, despite robust expression of the murine transgene as confirmed by qPCR (Fig. S3D). Because DPYSL2 promotes STAT3 activation through interaction with JAK1, we next asked whether DHP depletion affects the formation of the DPYSL2-JAK1 complex. FLAG immunoprecipitation experiments revealed markedly reduced JAK1 co-immunoprecipitation with DPYSL2-FLAG in DPYD-knockout cells compared with vector control cells (Fig. 3G). Importantly, quantification of the JAK1/FLAG ratio demonstrated a significant reduction in JAK1 association (∼78%) after normalization to the amount of immunoprecipitated DPYSL2, indicating that the decrease in complex formation cannot be explained solely by reduced DPYSL2 abundance. These findings suggest that DHP depletion not only destabilizes DPYSL2 but also impairs its ability to interact with JAK1, providing a potential mechanistic basis for the loss of STAT3 signaling observed upon DHP depletion.

### The pseudoenzyme DPYSL2 binds DHU at a structurally conserved pocket

To investigate the structural basis of the interaction between DPYSL2 and DHU, we modeled the structures of DPYS and DPYSL2 bound to DHU and Zn ions using AlphaFold3 ^5151^(see Methods). Structural superposition of DPYS and DPYSL2 revealed strong global similarity (RMSD 0.47 Å) (Fig. 4A), alongside marked differences in the DHU-binding pocket. Several catalytic and substrate-positioning residues present in DPYS are substituted in DPYSL2, replacing charged or polar residues with nonpolar counterparts, thereby altering the pocket’s chemical features (Fig. 4A, enlarged). In DPYS, DHU occupies a catalytic pocket containing two Zn ions coordinated by H67, H69, K159, H192, and D326, together with substrate-positioning residues Y164 and N347 (Fig. 4B) with high confidence (ipLDDT: 89.17, iPAE: 3.15Å). Most of these residues are substituted in DPYSL2. Interestingly, in DPYSL2, these residues interact with DHU through a *single* Zn ion, coordinated by H198. Despite these changes, DHU is modeled in the corresponding pocket of DPYSL2 with high confidence (ipLDDT: 84.38, iPAE: 5.05Å). The model demonstrates that DHU is stabilized in the pocket through stacking-like packing involving R75 and H107, together with Zn-associated contacts. These changes generate a distinct interaction network with DHU, consistent with a remodeled binding interface that supports ligand stabilization rather than catalysis.

**Figure 4.**
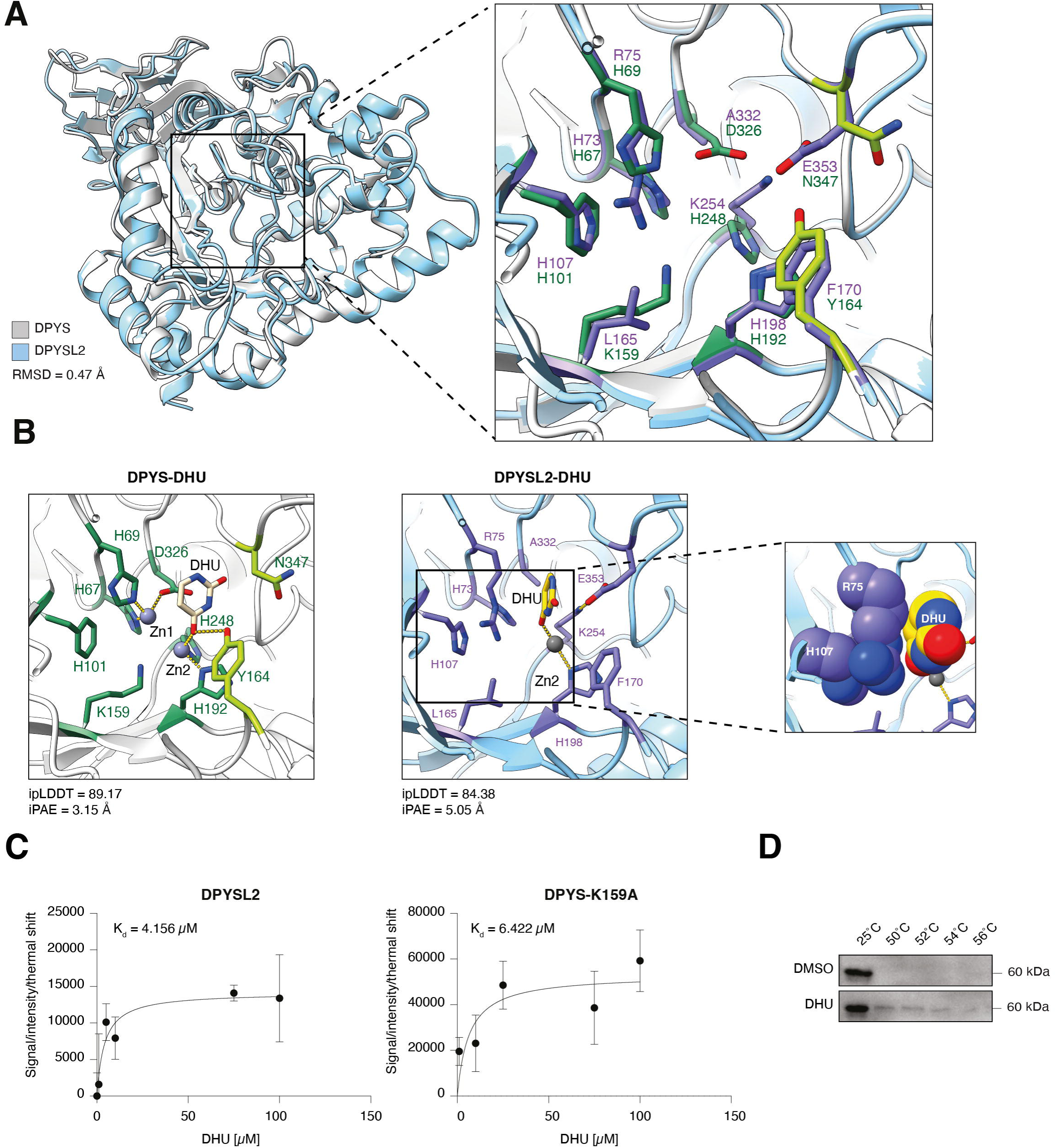
DPYSL2 contains a pseudoenzyme-like DHU-binding pocket and directly binds dihydrouracil. **(A)** DPYSL2 (sky blue) and DPYS (gray) show strong overall structural homology (RMSD = 0.47 Å), despite their different amino acid composition in the DHU-binding pocket (boxed region). In the magnified view of the binding pocket on the right, DPYS residues are depicted as green sticks (dark green, Zn-coordinating residues; light green, substrate-positioning residues), and the corresponding residues in DPYSL2 are shown as purple sticks. **(B)** In DPYS (left), DHU is captured by two Zn ions in the catalytic center (coordinated by H67, H69, K159, H192, H248, and D326) and substrate-positioning residues (Y164 and N347). In DPYSL2 (right), DHU is positioned within the corresponding pocket despite the absence of one Zn ion and the complete catalytic machinery. The DPYSL2 interface features Zn-associated contacts with a conserved His, an E353-K254 salt bridge, and a stacking-like packing with R75 and H107 (middle zoom-in, spheres). Both models are modeled with high interface confidence. Images were generated using ChimeraX v1.8. (**C**) Purified recombinant DPYSL2 or DPYS-K159A proteins were incubated with increasing concentrations of DHU and subjected to thermal shift analysis. DHU-induced changes in thermal stability were quantified and fitted using a one-site binding model to estimate the apparent dissociation constant (*K*d). DPYSL2 exhibited an apparent *K*d of 4.156 µM, whereas DPYS-K159A exhibited an apparent *K*d of 6.422 µM, indicating that DPYSL2 retains a DHU-binding affinity comparable to that of its enzymatic homolog. **(D)** DHU stabilizes recombinant DPYSL2 protein. Thermal stability assay of purified recombinant DPYSL2 in the presence of DMSO or DHU. Samples were heated to the indicated temperatures and subjected to immunoblotting with an antibody against DPYSL2.

To biochemically validate DHU binding, we performed an isothermal dose-response thermal shift assay (ITDR-TSA) on purified recombinant DPYSL2. DHU induced concentration-dependent thermal stabilization of the DPYSL2, demonstrating a direct interaction and allowing estimation of an apparent binding affinity (*K*d ≈ 4.2 µM) (Fig. 4C). To determine whether the predicted binding affinity was comparable to that of the DHP-binding protein DPYS, we performed the same analysis using recombinant DPYS-K159A, a catalytically impaired mutant that retains the predicted substrate-binding pocket. DPYS-K159A exhibited a similar apparent affinity for DHU (*K*d ≈ 6.4 µM), confirming the model prediction, in which DPYSL2 retains a DHP-binding capability comparable to that of its enzymatic homolog (Fig. 4C). Importantly, DHU failed to induce a measurable thermal shift in the unrelated protein LXXLL/leucine-zipper-containing ARF-binding protein (LZAP/CDK5RAP3), indicating that the observed stabilization is a specific property of DHU (Fig. S4A). Moreover, uracil, the pyrimidine precursor of DHU, did not induce thermal stabilization of either DPYSL2 or DPYS-K159A (Fig. S4B), demonstrating that the interaction is selective for the reduced pyrimidine metabolite DHU rather than pyrimidines in general. To independently validate this interaction, we separately performed a Thermal Shift Assay coupled with Western Blotting (TSA-WB), which confirmed DHU-dependent stabilization of DPYSL2 across a temperature gradient (Fig. 4D). Together, these results provide direct biochemical evidence that DPYSL2 directly interacts with DHU through a remodeled pocket, supporting a mechanism in which metabolite binding stabilizes DPYSL2 and promotes downstream STAT3 signaling.

### DHP depletion suppresses mesenchymal cell behavior and metastatic colonization

To determine whether DHP-dependent STAT3 signaling contributes to the maintenance of aggressive mesenchymal cell states, we examined the transcriptional, phenotypic, and functional consequences of DHP depletion. Transcriptomic analysis of DPYS-overexpressing MDA-MB-231 cells revealed significant downregulation of pathways linked to cell-cell adhesion, leukocyte activation, and matrisome-associated programs, indicating broad suppression of mesenchymal and extracellular matrix remodeling networks (Fig. 5A). Additionally, quantitative proteomic analysis revealed a coordinated reduction in extracellular matrix and adhesion-associated proteins, including ITGA6, LOXL2, LAMB3, ITGB4, SERPINE1, and MMP14 (Fig. 5B and Fig. S5A), supporting the collapse of the invasive mesenchymal proteome following DHP depletion.

**Figure 5.**
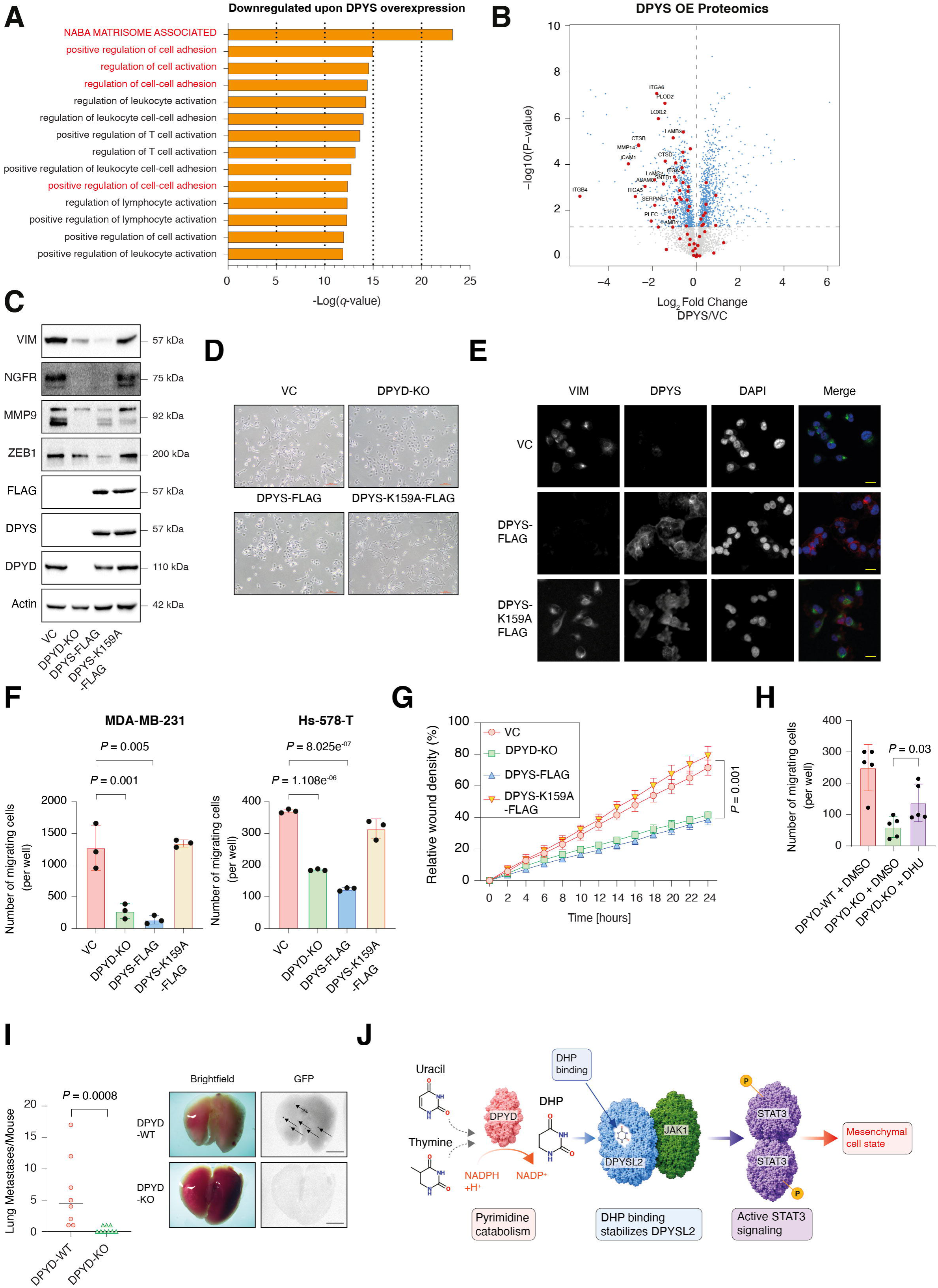
DHP depletion suppresses mesenchymal cell behavior and metastatic colonization. **(A)** DPYS overexpression suppresses adhesion- and matrisome-associated transcriptional programs. Vector control (VC) and DPYS-FLAG-overexpressing MDA-MB-231 cells were subjected to RNA-seq analysis. Genes significantly downregulated following DPYS overexpression were analyzed for pathway enrichment. Selected pathways and their corresponding *q* values are shown; the *q* values were calculated using the enrichment analysis tool. **(B)** DPYS overexpression suppresses ECM- and adhesion-associated proteins. Proteomic volcano plot comparing DPYS-FLAG-overexpressing MDA-MB-231 cells with vector control cells. Selected ECM- and adhesion-associated proteins are highlighted. **(C)** DHP depletion reduces the expression of mesenchymal and invasion-associated proteins. Immunoblots representing vector control (VC), DPYD knockout (DPYD-KO), DPYS-FLAG overexpression, and DPYS-K159A-FLAG expression in MDA-MB-231 cells. Cells were lysed and subjected to immunoblotting using the indicated antibodies. **(D)** DHP depletion alters mesenchymal cell morphology. Representative brightfield images of MDA-MB-231 cells expressing VC, DPYD-KO, DPYS-FLAG, or DPYS-K159A-FLAG. DPYD loss and DPYS overexpression result in a more compact morphology compared with VC and DPYS-K159A-FLAG cells. Scale bar, 100 µm. **(E)** DPYS catalytic activity is required to disrupt the organization of mesenchymal VIM filaments. VC, DPYS-FLAG-overexpressing, and DPYS-K159A-FLAG-expressing MDA-MB-231 cells were subjected to immunofluorescence imaging using the indicated antibodies. Scale bar, 10 µm. **(F)** DHP depletion suppresses migration in basal breast cancer cell lines. Transwell migration assays in MDA-MB-231 and Hs-578-T cells expressing VC, DPYD-KO, DPYS-FLAG, or DPYS-K159A-FLAG. Quantification is reported as the number of migrating cells per well. Each value represents the mean ± s. d. (*n* = 3). The *P* value was determined by an unpaired two-sided Student’s t-test. **(G)** DHP depletion impairs wound closure. Real-time quantification of relative wound density in the indicated MDA-MB-231 cell lines using the Incucyte Live-Cell Analysis System. Cells were monitored over 24 h. Each value represents the mean ± s.d. (*n* = 8). The *P* value was determined by the Wilcoxon rank-sum test. **(H)** DHU supplementation partially rescues migration in DPYD-KO cells. Transwell migration assay of DPYD-WT cells treated with DMSO and DPYD-KO cells treated with either DMSO or 100 µM DHU. Quantification is reported as the number of migrating cells per well. Each value represents the mean ± s.d. (*n* = 5). The *P* value was determined by the Wilcoxon rank-sum test. **(I)** DPYD loss suppresses lung metastatic colonization. GFP-labeled VC or DPYD-KO MDA-MB-231 cells were injected into the tail vein. Left: representative brightfield and GFP images of lungs. Right: quantification of lung metastases per mouse. Each value represents one mouse. The *P* value was determined by the Wilcoxon rank-sum test. **(J)** Model for DHP-mediated regulation of DPYSL2 stability and STAT3 signaling. Schematic model illustrating how DPYD-generated DHPs bind and stabilize DPYSL2, thereby sustaining JAK1-STAT3 signaling and promoting mesenchymal cell states, migration, and metastatic colonization. Depletion of intracellular DHPs destabilizes DPYSL2, attenuating STAT3 signaling and suppressing mesenchymal and metastatic phenotypes.

We next validated these pathway-level observations at the molecular level. Immunoblot analysis confirmed that DHP depletion reduced the abundance of canonical mesenchymal and invasion-associated proteins, including VIM, ZEB1, MMP9, and NGFR (Fig. 5C). Quantitative RT-PCR analysis similarly demonstrated downregulation of multiple mesenchymal, inflammatory, and extracellular matrix-associated genes, including *MMP14, MMP9, COL7A1, COL6A2, SNAI2, NGFR, L1CAM*, and *EFEMP2* (Fig. S5B).

Consistent with these molecular alterations, DHP depletion induced a striking change in cell morphology, shifting cells away from their distinct spindle-shaped mesenchymal appearance toward a more compact epithelial-like morphology (Fig. 5D). This phenotype was further reinforced by immunofluorescence analysis, where VIM filament organization was markedly disrupted in DPYS-overexpressing cells but remained largely preserved in cells expressing DPYS-K159A, demonstrating that relatively high DHP levels are required to maintain mesenchymal cytoskeletal architecture (Fig. 5E). Similarly, immunofluorescence staining for EZRIN revealed a pronounced loss of lamellipodial structures upon DPYD loss, further supporting disruption of mesenchymal cell organization following DHP depletion (Fig. S5C).

We next asked whether these molecular and morphological changes translate into impaired migratory capacity. In both MDA-MB-231 and Hs-578-T basal B breast cancer cell lines, DHP depletion significantly reduced transwell migration (Fig. 5F and Fig. S5D). Utilizing the Incucyte Live-Cell Analysis System, wound-healing assays further demonstrated a marked reduction in relative wound density recovery following DPYD loss or DPYS overexpression, whereas the K159A mutant remained largely comparable to vector control cells (Fig. 5G and S5E). Moreover, we found that DHU supplementation significantly increased intracellular DHU abundance (Fig. S5F) and partially rescued the migration defect of DPYD-deficient cells (Fig. 5H and Fig. S5G). Additionally, we found that restoration of DPYD expression significantly rescued the migratory defect associated with DPYD loss (Fig. S5H), further establishing that DPYD-dependent DHP production is required to sustain the migratory capacity of mesenchymal breast cancer cells. Finally, extending these findings *in vivo*, tail-vein injection assays revealed a profound reduction in lung metastatic colonization following DPYD loss, establishing that intracellular DHP abundance is required not only for mesenchymal signaling and migration *in vitro*, but also for efficient metastatic seeding *in vivo* (Fig. 5I). Collectively, these findings identify DHPs as metabolic regulators of mesenchymal cell identity and demonstrate that DHP depletion suppresses cytoskeletal organization, migration, and metastatic colonization.

## Discussion

Recent studies have established that metabolites can function not only as intermediates of metabolic flux, but also as signaling and regulatory molecules ^19,20^. These metabolites directly influence cancer biology by reshaping epigenetic states, transcriptional programs, and signal transduction pathways, as exemplified by classical oncometabolites. Most metabolite-signaling paradigms described to date have centered on central carbon metabolism, amino acid metabolism, or lipid-derived metabolites ^52^. By contrast, the signaling potential of nucleotide catabolic intermediates has remained largely unexplored. We previously showed that mesenchymal carcinoma cells activate an incomplete pyrimidine degradation pathway characterized by elevated DPYD but low DPYS expression, thereby promoting intracellular accumulation of DHPs ^16^. Here, we identify pyrimidine catabolism as an unexpected regulator of oncogenic signaling and demonstrate that the DHPs function as signaling metabolites that sustain JAK1-STAT3 pathway activity through stabilization of the DPYS homolog DPYSL2 (Fig. 5J). Specifically, genetic depletion of DPYD or enforced expression of catalytically active DPYS reduced intracellular DHP abundance, destabilized DPYSL2, suppressed STAT3 signaling, and impaired metastatic colonization. By contrast, catalytically inactive DPYS mutants failed to phenocopy these effects, demonstrating that DHP levels, rather than enzyme expression per se, govern these cellular effects. Mechanistically, our study uncovers a previously unrecognized mode of metabolite-dependent signaling regulation. Whereas many signaling metabolites function through covalent chromatin modifications or enzymatic cofactor activity ^20^, DHPs appear to regulate signaling through direct stabilization of a scaffold-like signaling protein. DPYSL2 belongs to the collapsin response mediator protein (CRMP) family ^53^ and shares a strong structural homology with the metabolic enzyme DPYS, despite lacking catalytic activity. Our findings raise the intriguing possibility that DPYSL2 has evolutionarily retained the capacity to bind pyrimidine catabolites, repurposing this interaction for signaling regulation rather than metabolism. In this framework, DHP binding may stabilize DPYSL2 conformation or protect it from proteasomal degradation, thereby sustaining the assembly or activity of signaling complexes required for efficient STAT3 activation.

The coupling between pyrimidine catabolism and STAT3 signaling is particularly important in mesenchymal and metastatic cancer cell states. STAT3 functions as a central regulator of EMT, inflammatory signaling, metastatic dissemination, and therapy resistance across multiple tumor types. Our analyses demonstrated that DPYD expression strongly co-segregated with EMT-associated transcriptional programs, inflammatory signaling pathways, and aggressive tumor subtypes across diverse cancers. These observations suggest that the accumulation of DHPs contributes to maintaining mesenchymal cell identity by sustaining the signaling cues required to sustain STAT3 activity. More broadly, our findings support the emerging concept that metabolic states can stabilize cell-state programs not only by supplying biosynthetic intermediates but also by directly reinforcing signaling networks that govern cellular plasticity.

An important implication of this work is that pyrimidine catabolism regulates signaling beyond cancer cells. DPYD expression is particularly elevated in myeloid cells and tumor-associated macrophages, where it has been implicated in fluoropyrimidine metabolism and immune regulation ^54^. Given the central role of STAT3 signaling in macrophage polarization, inflammatory suppression, and tumor immune evasion, it is tempting to speculate that DHP-mediated signaling operates within the tumor immune microenvironment. Although our current study focuses on cancer-cell-intrinsic functions of pyrimidine catabolism, future work will be required to determine the involvement of DHPs in immune-cancer cell interactions.

Our findings have therapeutic implications as pharmacological targeting of DPYD has primarily been explored in the context of fluoropyrimidine metabolism and 5-FU toxicity. Our results suggest that inhibition of pyrimidine catabolism further suppresses aggressive mesenchymal tumor states by destabilizing DPYSL2 and attenuating STAT3 signaling. Beyond the specific context of pyrimidine metabolism, the identification of a metabolite-dependent protein stabilization mechanism suggests that targeting metabolite-protein interactions is a tractable strategy to disrupt oncogenic signaling states maintained by metabolic rewiring.

---

**Figure S1. DPYD expression is associated with mesenchymal and inflammatory signaling programs across cancers (A)** DPYD expression positively correlates with EMT and inflammatory signaling pathways across multiple cancer types. Gene set enrichment analysis using the GENI platform shows Hallmark pathways positively associated with *DPYD* expression in glioblastoma, lung cancer, and pancreatic cancer datasets (TCGA, Firehose Legacy). Pathways are ranked by FWER *P* value, and point color indicates FDR *q* value. Statistical significance was assessed using the GENI GSEA workflow, with FWER *P* values and FDR *q* values shown. **(B)** DPYD expression correlates with EMT-associated transcriptional programs. GSEA plots showing enrichment of the Hallmark EMT pathway in gene expression profiles ranked by their correlation with *DPYD* across breast, lung, pancreatic cancer, and glioblastoma datasets (TCGA, Firehose Legacy). Normalized enrichment scores (NES), FWER *P* values, and FDR *q* values are indicated. **(C)** TGFβ-induced EMT increases DPYD expression. A549 lung adenocarcinoma cells were treated with TGFβ for the indicated time points, lysed, and subjected to immunoblotting using the indicated antibodies. **(D)** Basal breast cancer cell lines express elevated DPYD levels and low DPYS expression. Luminal and basal breast cancer cell lines, together with MDA-MB-231 cells ectopically expressing DPYS, were lysed and subjected to immunoblotting using the indicated antibodies. **(E)** Mesenchymal breast cancer cell lines selectively express DPYD but lack DPYS expression. Luminal and basal breast cancer cell lines and normal mouse liver tissue were lysed and subjected to immunoblotting with the indicated antibodies.

**Figure S2. Validation of DHP manipulation, DPYS catalytic activity, and STAT3 transcriptomic suppression (A)** Conservation of catalytic and substrate-binding residues across structurally characterized DPYS homologs. Multiple sequence alignment of human DPYS with structurally characterized dihydropyrimidinase (DPYS) homologs from *Saccharomyces cerevisiae* and *Dictyostelium discoideum*. Conserved amino acids are highlighted according to sequence similarity. Residues implicated in Zn coordination are marked with green asterisks, whereas residues involved in DHU binding are indicated by orange asterisks. The catalytic lysine residue (K159) in human DPYS is highlighted and is conserved among the aligned homologs. **(B)** The DPYS-K159L mutant lacks DHU catabolic activity. *In vitro* DPYS enzymatic activity assay measuring DHU turnover by wild-type DPYS and DPYS-K159L. DHU absorbance was measured at the indicated time points. **(C)** The uracil-derived branch predominates over the thymine-derived branch in MDA-MB-231 cells. Targeted metabolite profiling of uracil, DHU, thymine, and DHT in MDA-MB-231 cells. Metabolite concentration is presented on a log10 scale. DHT levels remained below the limit of quantification. Each value represents the mean ± s.d. (*n* = 3). **(D)** Manipulation of DHP abundance does not significantly impact cell proliferation. MDA-MB-231 and Hs-578-T cells expressing vector control (VC), DPYD knockout (DPYD-KO), DPYS-FLAG, or DPYS-K159A-FLAG were monitored for proliferation. Each value represents the mean ± s.d. (*n* = 6). The *P* value was determined by one-way ANOVA. **(E)** DPYS overexpression induces a major global transcriptional shift. Principal component analysis of RNA-seq data from vector control and DPYS-FLAG-expressing MDA-MB-231 cells. **(F)** DPYS overexpression remodels the secretome. Volcano plot showing differential protein abundance following DPYS overexpression in MDA-MB-231 cells. Secreted and extracellular matrix-associated proteins are highlighted. **(G)** DPYS overexpression suppresses multiple signaling pathways. A summary dot plot of significantly downregulated gene sets identified by GSEA in DPYS-FLAG-expressing MDA-MB-231 cells relative to vector control cells. Gene sets are ranked by the FWER *P* value. Dot size represents gene set size, and dot color indicates FDR *q* value. **(H)** IL6-JAK-STAT3 signaling-associated genes in vector control and DPYS-FLAG-expressing MDA-MB-231 cells. (**I**) DPYS overexpression suppresses STAT3-associated transcriptional programs. Volcano plots showing the expression of DoRothEA-defined STAT3 target genes. (**J**) RNA-seq findings were validated by quantitative RT-PCR. Quantitative RT-PCR analysis of the indicated genes in vector control (VC), DPYD knockout (DPYD-KO), DPYS-FLAG overexpressing, and DPYS-K159A-FLAG expressing MDA-MB-231 cells. Each value represents the mean ± s.d. (*n* = 3). **(K)** Doxycycline induces robust expression of DPYS-GFP and DPYS-K159A-GFP. Representative fluorescence microscopy images of doxycycline-inducible DPYS-GFP and DPYS-K159A-GFP MDA-MB-231 cells at the indicated time points. Scale bar, 50 µm.

**Figure S3. DPYSL2 regulation by DHPs occurs primarily at the protein level (A)** DPYD loss reduces DPYSL2 protein levels at the single-cell level. DPYD-WT and DPYD-KO MDA-MB-231 cells were subjected to immunofluorescence imaging using the indicated antibodies to visualize DPYSL2 and DPYD at the single-cell level. DAPI marks nuclei. Scale bar, 10 µm. **(B)** Quantitative RT-PCR analysis of DPYD, DPYS, and DPYSL2 mRNA levels in vector control (VC), DPYD-KO, DPYS-FLAG, and DPYS-K159A-FLAG MDA-MB-231 cells. Each value represents the mean ± s.d. (*n* = 3). **(C)** DPYSL2 exhibits extensive sequence similarity to DPYS while lacking key catalytic features. Amino acid sequence alignment of human DPYS and DPYSL2. Conserved residues are highlighted. The catalytic lysine residue in DPYS (K159), which is required for DHP hydrolysis, and the corresponding region in DPYSL2 are indicated. The alignment demonstrates extensive sequence conservation between DPYS and DPYSL2 despite the lack of DPYS enzymatic activity in DPYSL2. **(D)** Quantitative RT-PCR analysis of DPYSL2 and murine Dpyd (mDpyd) mRNA levels in DPYD-KO MDA-MB-231 cells reconstituted with wild-type murine Dpyd (mDpyd) or the catalytically attenuated mDpyd-I560S mutant. Each value represents the mean ± s.d. (*n* = 3).

**Figure S4. DHU binding to DPYSL2 is selective and not a nonspecific property of proteins.** (**A**) DHU does not induce thermal stabilization of LZAP. Thermal shift analysis of recombinant LZAP incubated with increasing concentrations of DHU. Unlike DPYSL2 and DPYS-K159A, LZAP did not exhibit a measurable DHU-dependent thermal shift, indicating that DHU does not nonspecifically bind or stabilize unrelated proteins. (**B**) Uracil does not bind DPYSL2 or DPYS-K159A. Thermal shift analysis of recombinant DPYSL2 (left) and DPYS-K159A (right) incubated with increasing concentrations of uracil. No concentration-dependent thermal stabilization was observed, indicating that binding is selective for DHU rather than pyrimidines in general.

**Figure S5. DHP depletion suppresses extracellular matrix remodeling and mesenchymal cell behavior (A)** DPYS overexpression suppresses extracellular matrix organization proteins. Proteomic analysis comparing vector control (VC) and DPYS-overexpressing MDA-MB-231 cells. Proteins belonging to the REACTOME_EXTRACELLULAR_MATRIX_ORGANIZATION gene set were compared with all other detected proteins. The distribution of log2 fold-change values is shown. The *P* value was determined by the Wilcoxon rank-sum test. **(B)** DHP depletion suppresses mesenchymal and invasion-associated gene expression. Quantitative RT-PCR analysis of the indicated genes in vector control (VC), DPYD knockout (DPYD-KO), DPYS-FLAG overexpressing, and DPYS-K159A-FLAG expressing MDA-MB-231 cells. Each value represents the mean ± s.d. (*n* = 3). **(C)** DPYD loss alters cytoskeletal organization. VC and DPYD-KO MDA-MB-231 cells were subjected to immunofluorescence imaging using the indicated antibodies to visualize EZRIN and DPYD at the single-cell level. Scale bar, 10 µm. **(D)** DHP depletion suppresses migration in basal breast cancer cell lines. Representative images of transwell migration assays performed in MDA-MB-231 and Hs-578-T cells expressing vector control (VC), DPYD-KO, DPYS-FLAG, or DPYS-K159A-FLAG. Scale bar, 100 µm. (**E**) DHP depletion impairs wound closure. Representative images from wound-healing assays performed in MDA-MB-231 cells expressing VC, DPYD-KO, DPYS-FLAG, or DPYS-K159A-FLAG. Images were acquired at 0 and 24 h following scratch induction. **(F)** Exogenous DHU enters DPYD-KO cells. Targeted metabolite analysis of intracellular DHU levels following 100 µM DHU supplementation in DPYD-KO cells. Each value represents the mean ± s.d. (*n* = 3). The *P* value was determined by an unpaired two-sided Student’s t-test. **(G)** DHU supplementation partially restores migration in DPYD-KO cells. Representative images of transwell migration assays performed in DPYD-WT cells treated with DMSO and DPYD-KO cells treated with either DMSO or 100 µM DHU. Scale bar, 100 µm. (**H**) Reconstitution of DPYD restores migratory capacity. Representative images (left) and quantification (right) of transwell migration assays performed in VC cells, DPYD-KO cells expressing an empty vector, or DPYD-KO cells reconstituted with wild-type murine Dpyd (mDpyd). Quantification is reported as the number of migrating cells per well. Each value represents the mean ± s.d. (*n* = 3). The *P* values were determined by the Wilcoxon rank-sum test.

## Methods

### Data analysis

Gene set enrichment analysis (GSEA) for single-gene and differential expression data was performed using the GENI platform (https://yoavshaul-lab.shinyapps.io/gsea-geni/). Enrichment analysis of the differential expression data was performed using Metascape ^55^. Patient genomic and proteomic data were retrieved from cBioPortal. Cell line genomic data were obtained from the MERAV platform. Single-cell RNA sequencing (scRNA-seq) data ^56^ and secretome gene sets ^11^ were obtained from previously published research ^11^. STAT3 target genes were defined using the DoRothEA platform ^47^. DoRothEA assigns a confidence level (A to E) to each interaction based on the number of lines of supporting evidence. To ensure data robustness, analyses were restricted to interactions with confidence levels A through D, thereby retaining only experimental or literature-curated evidence. Datasets were analyzed using R (version 4.4.1).

### Alignment and modeling

The amino acid sequences of dihydropyrimidinase (*Saccharomyces kluyveri* (Q9P903) and *Sinorhizobium meliloti* (Q0PQZ5)), DPYS (Q14117), LZAP (Q96JB5), and DPYSL2 (Q16555) were obtained from UniProt ^57^. Multiple sequence alignments were generated with Clustal Omega ^58^, and an image was generated with ProtParam. Modeling and ligand-binding analysis were performed using AlphaFold3 ^51^ via a local implementation (https://github.com/Kuhlman-Lab/alphafold3, c6d8e0a), with the MMseqs2 server for generating MSA, selecting the best model among 10 seeds, and visualized using ChimeraX v1.8 ^5859^. Model confidence was estimated using the AlphaFold3 pLDDT and pAE values, focusing on interface protein residues and ligand atoms (i.e., those within 4 Å of a receptor atom).

### Cell lines

The cell lines ZR-75-1, MDA-MB-468, MCF7, CAL51, Hs-578-T, and MDA-MB-231 were obtained from ATCC (Manassas, VA, USA) and cultured in RPMI 1640 (Sartorius, Göttingen, Germany, 01-104-1A) or DMEM (Sartorius, 01-055-1A) supplemented with 10% FBS (Biological Industries, Beit Haemek, Israel, F7524), 20 mM L-glutamine (Sartorius, 03-020-1a), and 1% penicillin/streptomycin (Sigma, St. Louis, MO, USA, P0781-100ML). For TGFβ stimulation, cells were incubated with 5% FBS DMEM for 48 h, followed by stimulation with 5 ng/ml TGFβ (Peprotech, Cranbury, NJ, USA, 100-21) in 10 mM citric acid (pH 3.0) for the indicated time. Every 2 d, the cells were supplemented with media containing TGFβ. For doxycycline treatment, cells were incubated with 2 µg/ml doxycycline (Sigma, D3072) in DMEM for 96 h; the medium was replaced with fresh doxycycline-supplemented DMEM after 48 h.

### Cell lysis and immunoblotting

Cells were rinsed twice with ice-cold PBS (Sartorius, 020231A) and lysed with radioimmunoprecipitation assay (RIPA) buffer (20 mM Tris-HCl pH 7.4, 137 mM NaCl, 10% glycerol (vol/vol), 1% Triton X-100, 0.5% (wt/vol) deoxycholate, 0.1% (wt/vol) SDS, and 2 mM EDTA pH 8.0) supplemented with one tablet of EDTA-free protease inhibitor (Sigma, 11873580001) and phosphatase inhibitor cocktail (A and B, 100×) (Bimake, 100000184). The lysates were cleared by centrifugation at 21,000 × *g* for 10 min at 4°C. For mouse samples, livers were harvested from 6-week-old mice, sliced into 2 mm pieces, and homogenized in RIPA lysis buffer. To achieve complete solubilization, the homogenate was incubated on ice for 30 min with gentle intermittent mixing and finally passed through a 21-gauge needle 5–10 times. The lysate was cleared by centrifugation at 21,000 × *g* for 20 min at 4°C. The protein concentration was determined by Bradford assay (Bio-Rad, Hercules, CA, USA, 5000006). Proteins were denatured by adding SDS sample buffer (5×) and heated for 10 min at 95°C, resolved by 10% SDS-PAGE, transferred onto a 0.45 µm polyvinylidene difluoride membrane (Mercury, Burlington, MA, USA, IPVH00010), and probed with the appropriate antibodies. The images were quantified using Image Lab v6.1.0 (Bio-Rad).

### Immunoprecipitation

Cells were rinsed twice with ice-cold PBS and lysed in ice-cold lysis buffer (50 mM HEPES-KOH, pH 7.4, 2 mM EDTA, 10 mM pyrophosphate, and 1% NP40 alternative) supplemented with 0.5 mM sodium orthovanadate, 16 mM sodium fluoride, and one tablet of EDTA-free protease inhibitors per 50 ml. The lysates were cleared by centrifugation at 16,000 × *g* for 10 min at 4°C. Flag M2 affinity resins (Sigma, A2220) were washed with lysis buffer three times. Subsequently, 30 µl of a 50% resin slurry was added to the cleared cell lysates, and the mixture was incubated with rotation for 3 h at 4°C. For immunoblotting analysis, the beads were washed with lysis buffer supplemented with 150 mM NaCl. FLAG-tagged proteins were eluted by incubation in elution buffer (50 mM HEPES-KOH, pH 7.4, 2 mM EDTA, 10 mM pyrophosphate, 150 mM NaCl, 1% NP40 alternative, and 100 µg/ml FLAG peptide (Sigma, F4799)) for 1 h at 30°C with gentle shaking. Immunoprecipitated proteins were denatured by adding SDS sample buffer (5×) to a final volume of 150 µl and heating for 5 min at 95°C, resolved by 10% SDS-PAGE, and analyzed by immunoblotting as described above.

### Antibodies

Antibodies were obtained from the following sources: rabbit-mAb β-Actin (CST, Danvers, MA, USA, #4970), rabbit-mAb DPYD (CST, #4654), rabbit DPYS (Abcam, Cambridge, UK, ab205039), mouse-mAb FLAG (CST, #8146), rabbit-mAb Phospho-Stat3 (Tyr705) (CST, #9145), mouse-mAb Stat3 (CST, #9139), rabbit-mAb E-Cadherin (CST, #3195), rabbit-mAb N-Cadherin (CST, #13116), rabbit-mAb ZEB1 (CST, #3396), rabbit-mAb MMP-9 (CST, #13667), rabbit-mAb NGFR (CST, #8238), rabbit-mAb Vimentin (CST, #5741), rabbit-mAb DPYSL2 (CST, #35672), mouse-mAb DPYD (Abcam, ab54797), rabbit DPYS (Abcam, ab121844), mouse-mAb Vimentin (Abcam, ab8978), rabbit Ezrin (CST, #3145); HRP-labeled Goat anti-mouse (115-035-003), HRP-labeled Goat anti-rabbit (111-035-144), donkey anti-rabbit-Rhodamine red X (711-295-152), donkey anti-rabbit-Alexa Fluor 488 (711-545-152), donkey anti-mouse-Cy3 (715-165-150), and donkey anti-mouse-Alexa Fluor 488 (715-545-150) secondary antibodies were obtained from Jackson ImmunoResearch (West Grove, PA, USA). Measurement of IL-6 levels was performed by sandwich ELISA using the human IL-6 mini ABTS ELISA development kit (Peprotech, 900-M16)

### Virus production

HEK-293T cells were co-transfected with the pLJC2 (Addgene, Watertown, MA, USA #87975) or pLJM5 (Addgene, #61614), VSV-G envelope plasmid, and Δvpr lentiviral plasmid using PEI Transfection Reagent (Polysciences, Warrington, PA, USA, 24765). The supernatant containing the virus was collected 48 h after transfection and centrifuged at 400 × *g* for 5 min to eliminate cells and debris.

### CRISPR-Cas9-mediated KO cell lines

CRISPR-Cas9-mediated genome editing was utilized to achieve gene knockout, using the pSpCas9 (BB)-2A-Puro (PX459) V2.0 (Addgene, #62988), which delivers the sgRNA and Cas9 on a single plasmid. Cells were transfected and then subjected to single-cell cloning in 96-well plates. Editing of the *DPYD* locus was confirmed by immunoblotting to assess protein levels. Primers for cloning are sgRNA-*DPYD* 5’-CACCGTGTGCTCAGTAAGGACTCGG-3’ and 5’-AAACCCGAGTCCTTACTGAGCACAC-3’.

### Fluorescence-Activated Cell Sorting

For FACS analysis, MDA-MB-231 DPYSL2 GFP^+^ and DPYD KO DPYSL2 GFP^+^ cells (3.0 × 10^5^) were seeded in 6-well tissue culture plates with 10% FBS DMEM. The following day, cells were detached with a detachment buffer (0.05% EDTA in PBS) for 5 min, harvested in 10% FBS DMEM, and washed twice with ice-cold PBS. Finally, the cells were filtered through a round-bottom test tube with a 40-µm cell-strainer cap (Sigma, 352235) and analyzed using the FlowJo software (Tree Star, Ashland, OR, USA).

### RNA preparation and quantitative PCR (qPCR) analysis

Total RNA was isolated from cells using the NucleoSpin RNA Kit (Macherey-Nagel, Düren, Germany, MAN-740955), and reverse transcription was performed using the qScript cDNA Synthesis kit (Quantabio, Beverly, MA, USA, 95047). The resulting cDNA was diluted in DNase-free water (1:10) before quantification. The mRNA transcription levels were measured using 2× qPCR BIO SyGreen, Blue Mix Hi-ROX (NEB, Ipswich, MA, USA, M3003E), and StepOnePlus (Applied Biosystems). All data were expressed as the ratio of the target gene mRNA expression level to *ACTB* (*ACTIN*). Primer sequences obtained from Integrated DNA Technologies (IDT, Coralville, IA, USA) are provided in Supplementary Table 3.

### Transwell migration assay

Migration assay was performed using the 8.0-µm Pore Polycarbonate Membrane Transwell chamber (Corning, Corning, NY, USA, CA-3422). Transwell inserts were hydrated with serum-free DMEM for 30 min. MDA-MB-231 (2.0 × 10^4^) or Hs-578-T (2.0 × 10^4^) cells were suspended in 350 µl of serum-free DMEM and seeded in the upper chamber. To the lower chamber of the 24-well plate, 750 μl of 10% FBS in DMEM was supplemented. The medium was discarded after 20 h. The non-migratory cells were removed with cotton-tipped swabs, and the lower surface of the insert was stained with 0.05% crystal violet with methanol. The cells were counted and captured under a Nikon Eclipse 80i microscope at 10× magnification. Where indicated, cells were pre- treated with 100 µM DHU (ChemCruz, Dallas, TX, USA, 504-07-4) dissolved in DMSO every 12 h for 4 d prior to seeding.

### Wound healing assay

Cells (4.0 × 10^4^ per well) were seeded in IncuCyte ImageLock 96-well cell culture microplates (Sartorius, BA-04857). After 18 h, the cell monolayer was scraped using a wound-maker mechanical device (Essen BioScience, Ann Arbor, MI, USA), washed with PBS, and examined under an inverted microscope. The wound area was monitored using the IncuCyte live-cell imaging system. Wound healing assay results were compiled from eight wells, each containing one scratch. At the 24-h time point, closure of the control scratch was observed.

### Proliferation assay

Cells (500 per well) were plated onto white 96-well plates (Greiner, Kremsmünster, Austria, 655098) in 100 µl of 10% FBS DMEM. Cell viability was assessed with Cell Titer-Glo (Promega, Madison, WI, USA, G7570) on 1, 3, 5, and 7 d after seeding, and luminescence was measured with Cytation 3 Multi-Mode Reader (BioTek).

### Animal studies

A mouse experiment was conducted under an institutional protocol (MD-21-16429-5) approved by the Hebrew University Institutional Animal Care and Use Committee. The Hebrew University is certified by the Association for Assessment and Accreditation of Laboratory Animal Care. Sample sizes were determined based on previously published similar models ^10^, and no formal predetermination using statistical power analysis was performed. Mice were randomly allocated to experimental groups. MDA-MB-231 WT GFP^+^ and DPYD-KO GFP^+^ cells were injected into the tail vein of the six-week-old female NOD-SCID mice (1.0 × 10^6^ cells per mouse). After 4 weeks, the lungs were harvested and observed under the SMZ18 Nikon Stereomicroscope. The images were converted to gray scale and inverted. The pictures were slightly adjusted (levels) using Adobe Photoshop. Investigators were blinded to group allocation during outcome assessment.

### Fluorescence microscopy

Cells (7.0 × 10^4^) were seeded onto Poly-D-lysine (Sigma, P0899)-coated glass coverslips in 12-well plates. After 24 h, the slides were rinsed twice with PBS containing calcium and magnesium, and fixed with ice-cold methanol for 5 min. Next, cells were rinsed three times with PBS and incubated with the blocking buffer (1% BSA in TBST and 0.3 M glycine for quenching) for 30 min, followed by primary antibodies for 1 h (DPYD (Abcam, ab54797) 1:40, DPYS (Abcam, ab121844) 1:500, DPYSL2 (#35672, CST) 1:800, Ezrin (#3145, CST)1:50 and Vimentin (Abcam, ab8978) 1:1000) at room temperature. Cells were rinsed three times with PBS, incubated with secondary antibodies (1:200 in 1% BSA in TBST) for 1 h at room temperature in the dark following incubation with DAPI (1:100, Sigma, 15733122,) for 10 min at room temperature in the dark. Slides were mounted on glass coverslips using Vectashield (Vector Laboratories, Newark, CA, USA# H-1000). The cells were imaged on a Nikon Eclipse Ni-U upright fluorescence microscope (Nikon, Tokyo, Japan) equipped with a motorized NI-CTLB control unit and a Lumencor SOLA SE 5-LCR-VB LED light engine for epifluorescence illumination. Images were acquired using a 40×/60× objective with immersion oil and a Nikon DS-Qi2 monochrome CMOS camera, controlled by NIS-Elements BR software (v4.60.00, 64-bit; Nikon). The microscope was configured with appropriate fluorescence filter sets, including GFP and Cy3 channels. Image acquisition parameters were kept constant within each experiment, and images were saved as 16-bit files. The pictures were slightly adjusted (levels) using Adobe Photoshop.

### Hyper-IL-6 treatment

For Hyper-IL-6 production, HEK-293T cells were transfected with the desired plasmid ^48^ using PEI. The following day, half of the medium was replaced with 30% FBS DMEM. The supernatant containing the Hyper-IL-6 was collected 72 h after transfection and centrifuged at 400 × *g* for 10 min to remove cells and debris. Hyper-IL-6 concentration was determined using a sandwich ELISA using the human IL-6 mini ABTS ELISA development kit (Peprotech, 900-T16K). For the treatment, 1.0 × 10^6^ cells were starved in 5% FBS-DMEM overnight. Subsequently, cells were pre-treated with or without 1 µM ruxolitinib (ORNAT, Rehovot, Israel, FOB-10-4511-1) in serum-free DMEM (0% FBS) for 1 h. Finally, cells were treated with 5 pg/ml Hyper IL-6 in 0% FBS DMEM for 1 h.

### ELISA

IL-6 levels were measured by sandwich ELISA using the human IL-6 mini ABTS ELISA development kit according to the manufacturer’s instructions. Briefly, cells (1.0 × 10^5^ per well) were seeded in a 6-well plate in triplicate. Conditioned media were collected and cleared by centrifugation to remove cellular debris. Microplates were coated overnight at room temperature with 0.5 µg/ml capture antibody. Following a 1 h blocking step with 1% BSA, diluted samples and IL-6 standards were added and incubated for 2 h at room temperature. Plates were then sequentially incubated with 1 µg/ml detection antibody for 2 h and an Avidin-HRP conjugate for 30 min at room temperature, with extensive washing between steps. Color development was induced using the ABTS substrate, and absorbance was measured with the Cytation 3 Multi-Mode Reader.

### RNA-seq

Total RNA was isolated from cells using the NucleoSpin RNA Kit (Macherey-Nagel, MAN-740955), and quality was determined by TapeStation analysis (Agilent). Libraries were prepared from 1000-2000 ng of total RNA using the KAPA Stranded mRNA-Seq Kit (Kapa Biosystems, Wilmington, MA, USA, KK8421) according to the manufacturer’s instructions. Libraries were pooled to 10 nM and sequenced on an Illumina (San Diego, CA, USA) NextSeq 500 (86 bp single-end reads) at the Core Facility of the Hebrew University Faculty of Medicine. BCL output files were converted to FASTQ using bcl2fastq (v2.20.0.422). Reads were quality-trimmed and aligned to the human reference genome (GRCh38, Ensembl release 99) using TopHat v2.1.1 ^60^. Gene counts were quantified using htseq-count v2.01 ^61^, and genes with < 10 total counts were excluded. Normalization and differential expression analysis were performed using DESeq2 v1.36.0 ^62^. Differentially expressed genes (DEGs) were defined by an adjusted *P* value < 0.1, a baseMean > 5, and a baseMean-dependent log2 fold change threshold: absolute log2 Fold Change > 5/sqrt (baseMean) + 0.3.

### Proteomics

Cells (2.0 × 10^6^) were seeded on a 10-cm plate for 48 h, washed twice with ice-cold PBS, trypsinized, washed twice with PBS, and resuspended in 300 µl of Tris 25 mM, 5% SDS. Samples were then sonicated at an amplitude of 50 for 30 s. Finally, samples were centrifuged at 10,000 × *g* for 10 min. LC MS analysis was performed using a Q Exactive-HF mass spectrometer (Thermo Fisher Scientific, Waltham, MA, USA) coupled on-line to an Ultimate 3000 Dionex (Thermo Fisher Scientific) UHPLC. Data were acquired using Xcalibur software (Thermo Fisher Scientific). Mass spectra data were processed using the MaxQuant computational platform, version 2.6.2.0. Relative protein quantification in MaxQuant was performed using the label-free quantification (LFQ) algorithm.

### Liquid chromatography-mass spectrometry

Cells (1.0 × 10^6^) were seeded 48 h prior to extraction. On the day of extraction, cell culture plates were placed on ice, and cells were washed twice with ice-cold 0.9% NaCl. Subsequently, plates were transferred to dry ice, and 1 ml of pre-chilled (-80°C) 80% LC-MS-grade methanol containing internal standards (IS; 5-azacytosine, Sigma, 852473, and 5-bromouracyl, Sigma, 855049) was added to each plate. Cells were mechanically scraped, collected into microcentrifuge tubes, and vigorously vortexed for 10 min at 4°C. Standard samples were prepared similarly from separate 10 cm plates using 2.5 ml of the extraction buffer and subsequently aliquoted. The extracts were evaporated to dryness under a stream of nitrogen gas and stored at -80°C. Prior to analysis, the dried pellets were resuspended in 50 µl of LC-MS-grade water and vortexed. The homogenates were cleared by centrifugation at 21,000 × *g* for 10 min at 4°C. Finally, 40 µl of the supernatant from each sample was transferred into 300 µl fixed insert vials (Thermo Scientific, 03-FISV). LC-MS/MS analyses for pyrimidine and dihydropyrimidine were conducted on a Sciex (Framingham, MA, USA) Triple Quad™ 5500 mass spectrometer coupled with a Shimadzu (Kyoto, Japan) UHPLC System. The chromatographic separations were performed in an XSelect HSS T3 column 3.5 μm (100 × 2.1 mm, Waters Corporation, Milford, MA, USA). Data acquisition was performed using Analyst 1.6.3 software, and data were analyzed using MultiQuant 2.1 software.

### Protein purification

The expression vector containing the GST-tagged construct was transformed into E. coli T7. Transformed cultures were incubated at 37°C, then induced with isopropyl β-D-thiogalactopyranoside (IPTG, Biolab, 16242352) to a final concentration of 300 µM. Post-induction, the cultures were incubated for 20 h at 16°C. Cells were harvested and resuspended in lysis buffer containing 20 mM sodium phosphate, pH 8.0, 150 mM NaCl, 5 mM β-mercaptoethanol (β-ME), and 10 mM phenylmethylsulfonyl fluoride (PMSF). Cells were disrupted by a microfluidizer. The lysate was clarified by ultracentrifugation at 68,900 × *g* for 1 h at 4°C. The supernatant was applied to a GST-affinity column, and the protein was purified using an ÄKTA pure™ (Marlborough, MA, USA) protein purification system.

### Isothermal dose-response thermal shift assay (ITDR-TSA) and thermal shift assay (TSA-WB)

For ITDR-TSA, 2 µM of purified recombinant proteins in TBS buffer (100 mM NaCl, 20 mM Tris-HCl, pH 7.5) were incubated with 20 µM ZnCl2 (Sigma, 208086) and the indicated concentration of DHU for 30 min. Subsequently, samples were placed in a 96-well PCR plate with SyproOrange (Sigma, S5692-50UL, 1:80). The intensity of different concentrations of DHU was collected using StepOnePlus (Applied Biosystems), selected at a single temperature at the highest intensity of 0 µM DHU, and Kd was determined by one-site-specific binding Nonlinear Regression using GraphPad Prism version 11.0.0 (GraphPad Software, Boston, MA, USA). For TSA followed by WB, 2 µM of DPYSL2 protein in TBS buffer (100 mM NaCl, 20 mM Tris-HCl, pH 7.5) was incubated with DMSO or 100 µM DHU for 15 min at 24°C with gentle agitation (300 rpm). Samples were subjected to a temperature gradient (40-60°C) for 3 min in a thermal cycler, followed by incubation at room temperature for 3 min. Finally, samples were cleared as described above and subjected to immunoblotting.

### Enzymatic activity assay

The purified DPYS and DPYSL2 (70 ng) were incubated with 1 mM ZnCl2 in 100 mM Tris-HCl buffer (pH 8.0) for 3 min at room temperature. Subsequently, 2 mM DHU in 100 mM Tris-HCl (pH 8.0) buffer was added, and the DHU level was determined by measuring optical density (OD) at 230 nm every 30min until the endpoint using a Cytation 3 Multi-Mode Reader.

### Statistical analysis

Statistical analyses were performed using GraphPad Prism version 11.0.0 and R. Data are presented as mean ± s.d. Unless otherwise specified in the figure legends, statistical significance was determined using the unpaired two-sided Student’s t-test. For experiments with small sample sizes (*n* = 3), formal normality testing was not feasible. In these instances, individual data points are displayed in the figures to allow visual assessment of data distribution. The investigators were not blinded to allocation during experiments and outcome assessment. Apart from 3 mice that died prior to the experimental endpoint, no other animals or data points were excluded from the analysis. The exact sample size (*n*), representing biologically independent *in vitro* replicates or individual mice, is indicated in the figure legends. *In vitro* experiments were performed with at least three independent biological replicates, and *in vivo* experiments included 10 mice per group, randomly assigned to experimental groups. No statistical methods were used to predetermine sample sizes, but our sample sizes are consistent with those generally employed in the field.

## Data availability

The processed RNA-seq and proteomics data are provided in the Supplementary Tables. The raw sequencing and mass spectrometry data are currently being prepared for deposition to public repositories.

## Acknowledgment

We thank the members of the Shaul laboratory. The Genomic Applications Laboratory of the Core Research Facility at The Faculty of Medicine, The Hebrew University of Jerusalem, Israel, performed the RNA-Seq data analysis.

## Author contributions

A.H. designed the study, performed immunoblotting, *in vitro* functional assays, *in vivo* experiments, and wrote and edited the manuscript. S.K.-G. and O.S.-F. performed the AlphaFold3 structural analyses. A.K. performed immunoblotting, qPCR, flow cytometry, and contributed to manuscript writing and editing. V.G. and S.B. performed *in vitro* experiments. I.M. conducted the single-cell analyses. S.L. and S.S. performed immunoblotting and qPCR experiments. Y.S. performed flow cytometry analyses. R.G. generated and validated the doxycycline-inducible system. B.S. performed the *in vivo* experiments. A.A.R. performed the immunoprecipitation experiments. A.K. performed the ELISA experiments. M.L. performed molecular cloning, assisted with *in vitro* experiments, and edited the manuscript. R.W. performed *in vitro* experiments and provided experimental guidance. Y.D.S. conceived and supervised the study and wrote and edited the manuscript.

