## Supplementary figures and images for "Dihydropyrimidines sustain aggressive cancer states by stabilizing DPYSL2"

### Figure S1

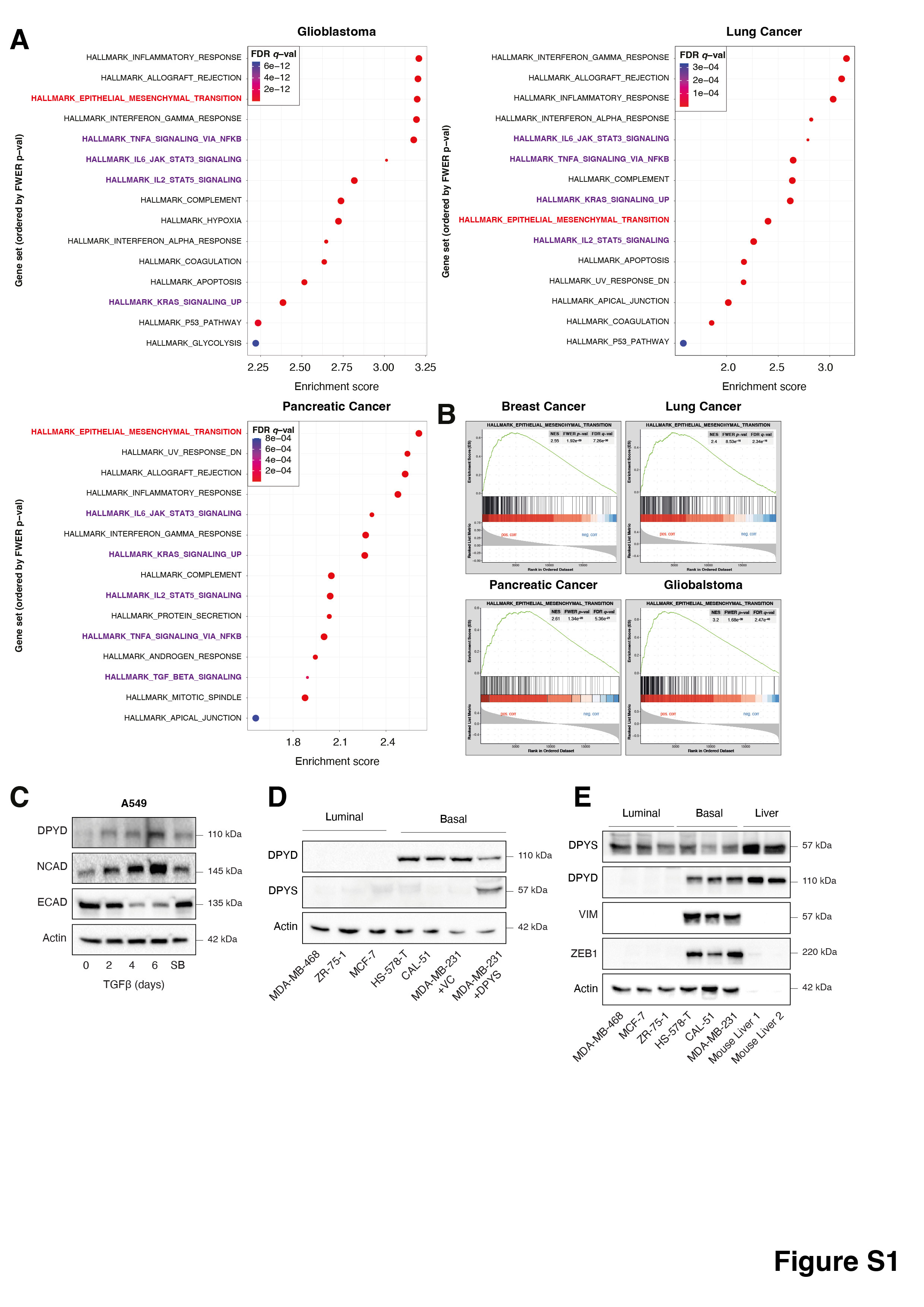

### Figure S2

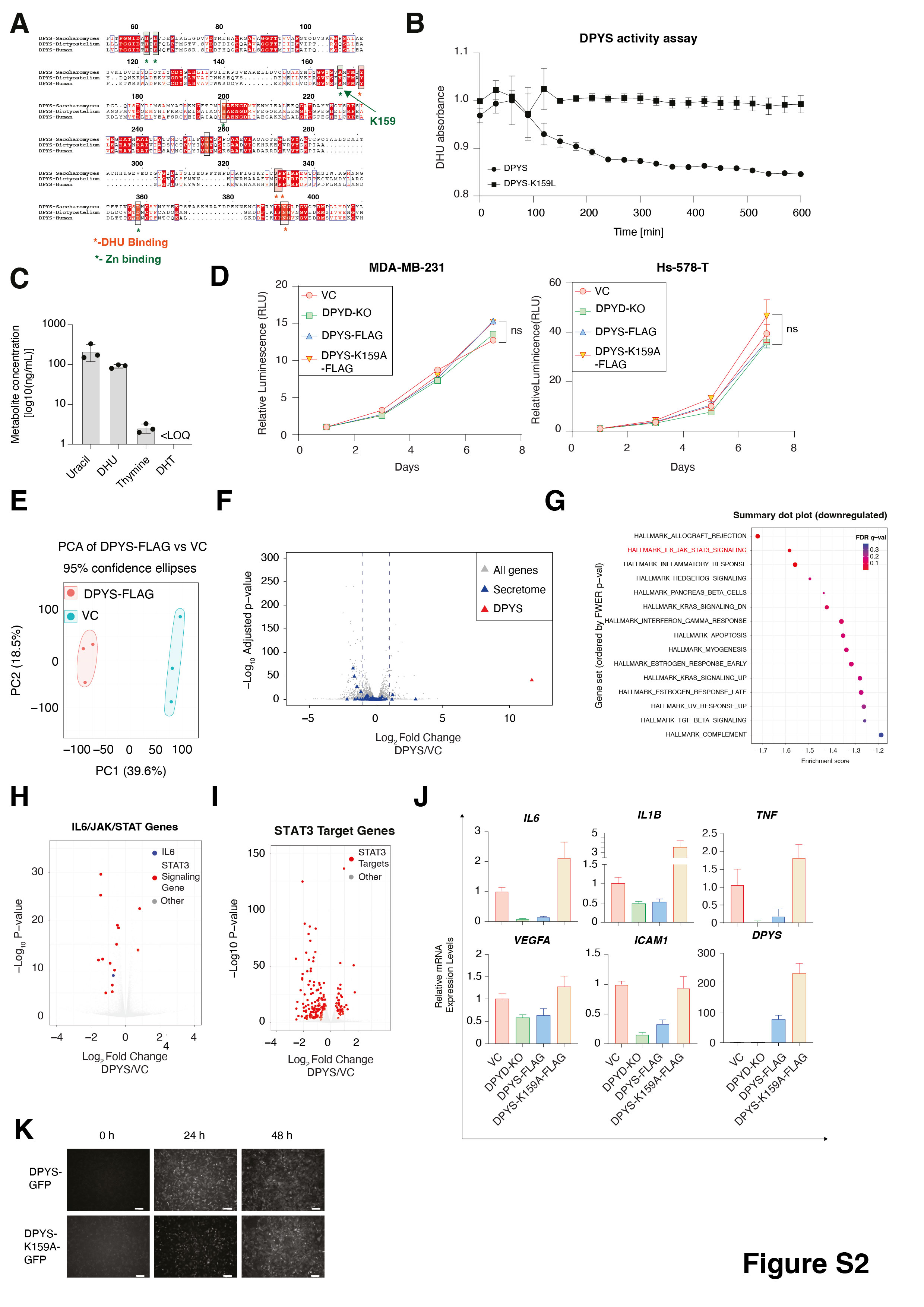

### Figure S3

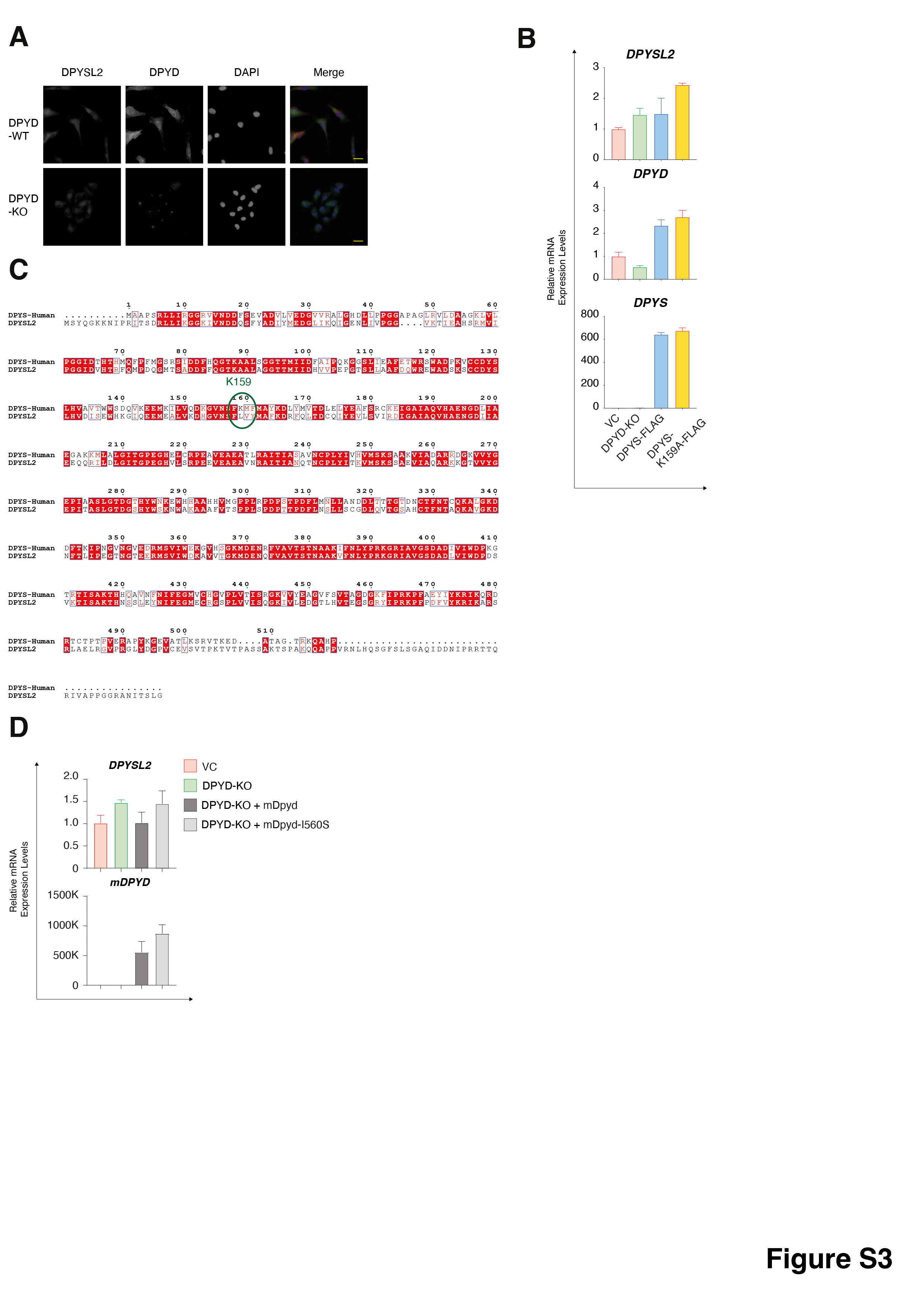

### Figure S4

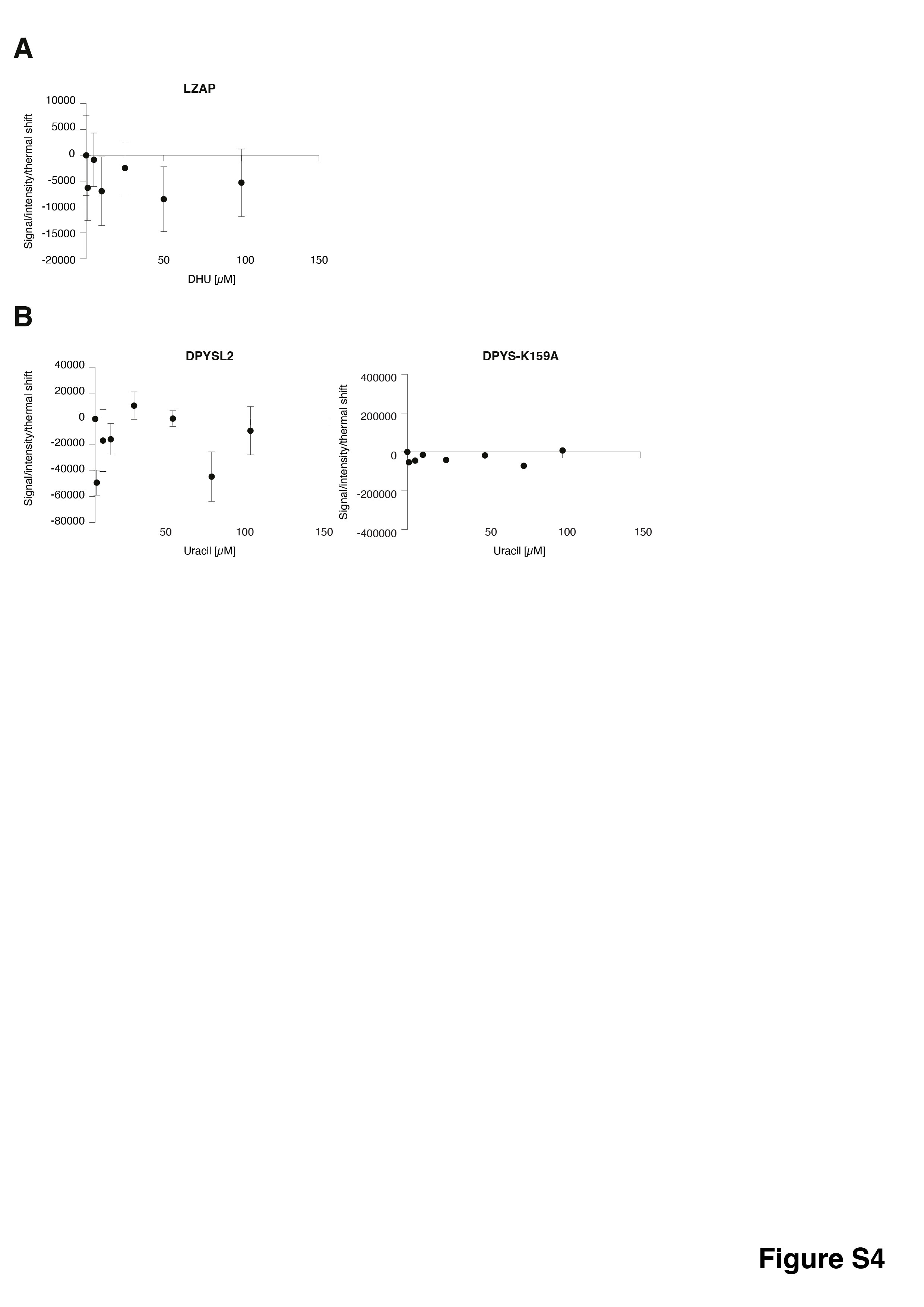

### Figure S5

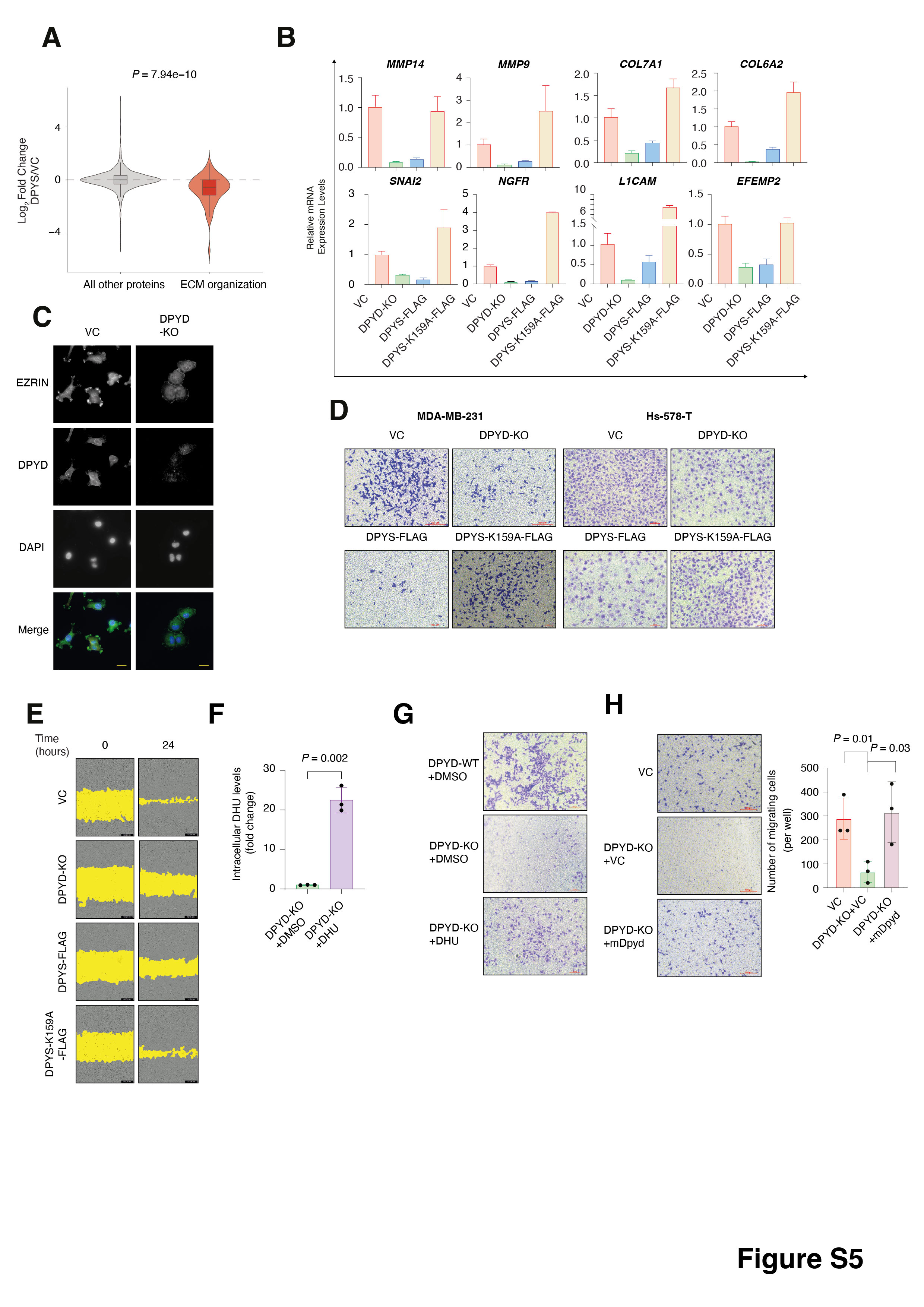
